# Disentangling abiotic and biotic effects of treated wastewater on stream biofilm resistomes enables the discovery of a new planctomycete beta-lactamase

**DOI:** 10.1101/2023.11.20.567610

**Authors:** Mustafa Attrah, Milo R. Schärer, Mauro Esposito, Giulia Gionchetta, Helmut Bürgmann, Piet N.L. Lens, Kathrin Fenner, Jack van de Vossenberg, Serina L. Robinson

## Abstract

**Background:** Antibiotic resistance, which is mediated by environmental reservoirs, poses a threat to human and animal health. Aquatic biofilms impacted by treated wastewater (WW) are known environmental reservoirs for antibiotic resistance, however the specific influence of biotic factors and abiotic factors from WW on the abundance of antibiotic resistance genes (ARGs) within aquatic biofilms remains unclear. Additionally, experimental evidence is limited as to whether genes with low sequence similarity to reference ARGs actually encode for functional ARGs, particularly within complex aquatic microbial communities.

**Results:** To disentangle the effects of abiotic and biotic factors on ARG abundances, natural biofilms were previously grown in flume systems with different proportions of stream water and either ultrafiltered or nonultrafiltered WW. In this study, we conducted deep shotgun metagenomic sequencing of 75 biofilm, stream, and WW samples from these flume systems and compared the taxonomic and functional microbiome and resistome composition. Statistical analysis revealed an alignment of the resistome and microbiome composition and a significant association with experimental treatment. Several ARG classes exhibited an increase in metagenomic abundances in biofilms grown with increasing percentages of nonultrafiltered WW. In contrast, sulfonamide and BEL family beta-lactamase ARGs showed greater abundances in biofilms grown in ultrafiltered WW compared to nonultrafiltered WW. Overall, our results pointed toward the dominance of biotic factors over abiotic factors in determining ARG abundances in WW-impacted stream biofilms and suggested gene family-specific mechanisms for ARGs which exhibited divergent abundance patterns. To investigate one of these specific ARG families experimentally, we biochemically characterized a new beta-lactamase from the *Planctomycetota* (*Phycisphaeraceae*). This beta-lactamase displayed activity in the cleavage of cephalosporin analog despite sharing low sequence identity with known ARGs.

**Conclusions:** This discovery of a functional planctomycete beta-lactamase ARG is noteworthy, not only because it was the first beta-lactamase to be biochemically characterized from this phylum, but also because it was not detected by standard homology-based ARG tools. In summary, this study conducted metagenomic analysis on the relative importance of biotic and abiotic factors in the context of WW discharge and their impact on both known and new ARGs in aquatic biofilms.

## Background

Antibiotic resistance is a major threat to public health. Among the numerous factors contributing to antibiotic resistance, anthropogenic pollution plays a role in altering the abundance and prevalence of antibiotic resistance genes (ARGs) in environmental reservoirs [1–4]. The rapid increase in the consumption of antibiotics and release of micropollutants (MPs) into the environment stimulates the spread of antibiotic-resistance genes (ARGs), posing a threat to modern medical procedures and treatments that rely on effective antibiotics [5].

Aquatic ecosystems that receive effluent from wastewater treatment plants (WWTPs) are influenced by biotic factors, such as antibiotic-resistant bacteria released from WWTPs [6] and abiotic factors [7,8] including nutrients and MPs that are not completely eliminated by conventional WWTPs [9]. Several recent studies have demonstrated that exposure to WWTP effluent alters the taxonomic and ARG compositions of microbial communities in downstream rivers [10–14]. However, disentangling the effects of abiotic and biotic factors on ARG abundance is challenging in the environment. Here, to systematically distinguish abiotic and biotic factors, we built on a previous research study in which controlled flume systems were used to grow biofilms in ultrafiltered and nonultrafiltered WWTP effluent diluted with different proportions of stream water [10,11]. Ultrafiltration (UF) treatment (pore size 0.4 µm) removed most of the microorganisms and particles (biotic factors) while maintaining consistent nutrient and MP levels, including antibiotics. Therefore, separating biotic factors from abiotic factors in UF treatment enabled investigation of the factors shaping the ‘resistome’, i.e., the total composition of antibiotic resistance genes (ARGs), of stream biofilms influenced by WWTP discharge.

Many studies on stream biofilms have relied on amplicon sequencing methods for taxonomic and/or ARG profiling [10,15–17]. In contrast to amplicon sequencing, short-read and long-read shotgun metagenomic sequencing studies of streams and stream biofilms [13,18,19] do not introduce polymerase chain reaction (PCR) amplification bias. As we demonstrated in this study, PCR-free metagenomic sequencing also enables the identification of sequence-divergent ARGs not detected by conventional primer pairs. Amplicon sequencing is more sensitive because it can aid in the detection of low abundance species, but it is also biased toward the amplification of specific taxa during PCR [20,21]. Specifically, PCR bias limits the detection of some bacterial phyla such as the *Planctomycetota*, for which the commonly-used 16S rRNA PCR primers do not properly bind [22,23]. The *Planctomycetota* are undersampled by classical amplicon-based methods [23] and are a focus in this study because of their relevance in WWTP [24], lacustrine [25], and riverine biofilm (periphyton) communities [26]. We conducted deep shotgun metagenomic sequencing of stream biofilms and water samples in this study, which enabled us to sample less well-studied microorganisms and their functional genes including ARGs.

Here, we demonstrated a combined computational and biochemical workflow to investigate the effects of treated wastewater (WW) on the microbiome taxonomic composition, functional enzyme diversity, and ARG abundance in flume biofilms grown in natural stream water mixed with different proportions of ultrafiltered and nonultrafiltered WW [10,11]. Specifically, we leveraged the functional genes identified through shotgun metagenomic sequencing to heterologously express and purify an uncharacterized ARG from the phylum *Planctomycetota*. Despite sharing low sequence identity with computationally-identified ARGs, the enzyme displayed beta-lactamase activity. Overall, this study enabled the dissection of biotic and abiotic factors which affect stream biofilm resistomes and led to the characterization of both known and new ARGs.

## Methods

### Biofilm cultivation and sampling

The flume systems and biofilm cultivation procedures used in two independent experiments to generate the samples sequenced in this study were described previously [10,11]. Briefly, the system consisted of flow-through channels fed with different mixtures of two different water sources: (1) a peri-urban stream (Chriesbach; Dübendorf, Switzerland) and (2) treated wastewater from a pilot-size wastewater treatment plant (100-person equivalents). In the first experiment (Exp. 1), the stream water was mixed with treated wastewater (WW) at the following percentages: 0% (control), 10%, 30%, and 80% WW. In the second experiment (Exp. 2), the 10% treatment was omitted and replaced by additional treatments with 30% and 80% ultrafiltered WW (UF) using a filter with a pore size of 0.4 µm to remove particles, e.g., suspended solids and microorganisms. Glass slides were added to the flow-through channels, and biofilms that naturally formed on the slides were allowed to develop for a period of four weeks. Water parameters (pH, temperature, conductivity, dissolved oxygen) were monitored weekly [10,11].

### DNA extraction and metagenomics assessment

Stream water, WW and UF WW, and biofilms from glass slides were sampled as described previously [11,27]. Total genomic DNA was extracted with a DNeasy PowerBiofilm Kit (QIAGEN) for a total of 75 samples (**Table S1**). The quality and quantity of the DNA extracts were measured using a NanoDrop One spectrophotometer. Extracted DNA samples were stored at -20°C and shotgun metagenomic sequencing was performed using the Illumina platform NovaSeq with a paired-end (2×150 bp; an average of 20 Gbp raw data per sample) strategy. Library construction and sequencing was performed by Novogene (Cambridge, UK). The reads obtained from shotgun metagenomics and those containing adapters and low-quality reads (N > 10% and quality score ≤ 5) were removed by Novogene, and the read qualities were double-checked by the authors using Fastp v.0.20.0 (https://github.com/OpenGene/fastp). De novo assembly was performed using MEGAHIT v1.2.9 [28]. Open reading frames (ORFs) were predicted from the assembled contigs using Prodigal v2.6.3 [29]. Although metagenomes from both Exp. 1 and Exp. 2 were sequenced and the data for both were deposited, we focused on Exp. 2 for the analyses reported here since Exp. 2 included UF treatments, unlike Exp. 1.

### Taxonomic and functional profiling

The taxonomic composition of the metagenomes was profiled using the degenerate consensus marker gene profiling tool, mTAGs v1.0.4, with 1000 maximum accepts and 1000 maximum rejects [30,31]. The mTAGs profiles for all samples were merged using mTAGs merge function. Genus- and phylum-level per-sample count tables were used for subsequent analysis. Counts corresponding to Enzyme Commission (EC) Number classes were compiled as described previously [32]. Briefly, fourth-level EC number counts were calculated using the EC-annotated prokaryotic fraction of the UniProt database using DIAMOND blastx v2.0.6 [33] and a minimum bitscore cutoff of 50 (all other parameters set to default).

### Reads-based ARG detection and ordination analyses

The ARG resistome was profiled using the DeepARG v1.0.2 short reads pipeline with default settings [34]. ARG counts normalized by the 16S-rRNA counts within the output files for each sample were merged into an abundance table. All data were analyzed and visualized in R v4.3.0 with the tidyverse package [35]. Principal coordinate analysis (PCoA) of the microbiome composition was performed using the genus abundance table. Unaligned and unassigned reads were removed. Genus counts were normalized using the trimmed mean of M-values (TMM) method within the EdgeR package [36–38]. Normalized genus counts were square-root-transformed. PCoA of the resistome composition was calculated from 16S-normalized DeepARG abundances [34]. PCoA analysis of functional enzyme composition was calculated from normalized and Hellinger-transformed EC numer counts. For all the data sets, pairwise Bray-Curtis distances between samples were computed using the ‘vegdist’ function from the vegan package [39]. PCoA was performed using the ‘pcoa’ function from the ape package [40]. Procrustes analysis comparing the PCoA vectors was conducted using the ‘procrust’ and ‘protest’ functions from the vegan package [39]. The Kruskal-Wallis test (McKnight and Najab, 2010) was applied to test for significant correlations between the normalized average coverage of different ARGs and wastewater effluent proportions (WW%), and p-values were corrected for multiple hypothesis testing using the Benjamini-Hochberg method [41].

### ARG identification and coverage analysis from metagenome assemblies

For a complementary and separate approach to the metagenomic short reads pipeline for ARG analysis, predicted open reading frames (ORFs) from assembled metagenomes were identified by protein BLAST v2.9.0 [42] search against the Structured Antibiotic Resistance Genes (SARG) protein database v2.2, with stringent e-values and query coverage cutoffs of 0.1 and 20, respectively [43]. Using thresholds established previously in similar samples [44] for the detection of ARGs in riverine metagenomes, BLAST hits with more than 85% amino acid identity and an alignment length of 100 amino acids were retained. BLAST hits were then clustered using CD-HIT v4.6.8 [45,46], with parameters –n 5 and –c 0.7. The 102 non-singleton cluster representatives of SARG hits were selected for further analysis. Metagenomic coverage of selected ARG cluster representatives were calculated across all metagenomes based on methods previously described [44]. Briefly, raw reads were mapped back to BLAST hits using Bowtie2 v2.4.4 [47] with the parameters shown in **Table S2**. Coverage depth was calculated using samtools v1.12 and average coverage was calculated as described previously [48]. The average coverage of the resulting 102 ORFs was normalized to reads per million as previously described [44].

### Bioinformatic analyses of beta-lactamases

Metallo-beta lactamase fold proteins were identified by querying assembled proteins from the 75 metagenomes and additional publicly-available metagenomes from MP enrichment studies (NCBI accession: PRJNA725625) using BLAST v.2.9.0 [49] with relaxed parameters (e-value = 0.1, qcov_hsp_perc = 20, all other parameters set to default) to detect homologs with low sequence identity to known ARGs. Taxonomic diversity and trends in abundances of metallo-beta lactamase fold homologs across different WW and UF WW conditions were examined leading to the prioritization of two metallo beta-lactamase homologs (NCBI accession numbers MBX3358097.1 and MBX3359331.1) for experimental characterization. Structural modeling of MBX3358097.1 was conducted using ColabFold AlphaFold 2.0 [50] with default parameters. AlphaFold models were assessed for quality based on predicted local distance difference test (pLDDT) scores and aligned pairwise with the reference structure in the Protein Data Bank (PDB ID: 3NKS) using the PyMOL ‘super’ command with default parameters.

### Cloning and protein expression

For functional characterization, two full-length *Phycisphaeraceae* (*Planctomycetota*) proteins (NCBI accession numbers MBX3358097.1 and MBX3359331.1) were codon-optimized for *E. coli* codon usage using the Integrated DNA Technologies Codon Optimization Tool (https://eu.idtdna.com/pages/tools/). Signal peptides were detected using SignalP v6.0 [51] and removed where present (**Table S3**). The genes were synthesized as gBlocks by Integrated DNA Technologies. The genes were cloned individually by Gibson Assembly into the multiple cloning site 1 of the pETDuet-1 vector (*ampR* resistance marker) with N-terminal hexahistidine tags (**Table S3**). The constructs were transformed into *E. coli* DH5α cells and verified by Sanger sequencing (Microsynth, Balgach, Switzerland). The sequence-verified constructs were subsequently transformed into *E. coli* BL21(DE3) cells (New England Biolabs Frankfurt, Germany). Starter cultures (5 mL) were grown in LB with 100 μg/ml ampicillin overnight at 37°C. From the starter cultures, 2.5 mL was added to 400 mL LB with 100 μg/mL ampicillin in a 1 L baffled Erlenmeyer flask and grown to an optical density of 0.5-0.8 at 37°C. Protein expression was induced by adding 0.1 mM isopropyl β-D-1-thiogalactopyranoside (IPTG) and cultures were incubated for 18 h at 16°C.

### Protein purification

Expression cultures were poured into 50 mL Falcon tubes and centrifuged at 4000 rcf for 15 min. Supernatants were decanted, and cell pellets were resuspended in 4 mL buffer A (20 mM Tris, 500 mM NaCl, 20 mM imidazole, 10% glycerol, pH = 8) per gram of pellet weight. Cells were lysed by 2 cycles of sonication using a 6 mm needle, 20% amplitude and 10 x 10 second pulses with a rest time of 10 seconds in between (total sonication time per cycle = 3 minutes). Lysates were centrifuged at 20,000 rcf at 4°C for 1h. Cleared lysate was filtered through a 0.2 μm syringe filter, loaded into a ÄKTA Pure system (Cytiva Europe GmbH, Glattbrugg, Switzerland) and injected onto a HisTrap FF 5 mL column (Cytiva, Europe GmbH, Glattbrugg, Switzerland). The column was washed with 20 column volumes of buffer A. His-tagged proteins were eluted in 3 column volumes of buffer B (same as buffer A but with an imidazole concentration of 500mM), and 8 × 2 mL fractions were collected. The fractions with the highest protein concentrations were determined qualitatively by Bradford assay [52]. Imidazole was removed by running the sample over a PD-10 desalting column (Cytiva Europe GmbH, Glattbrugg, Switzerland) and eluting in buffer (5 mM Tris, 30 mM NaCl, 10% glycerol, 0.1 mM ZnCl_2,_ pH = 8). ZnCl_2_ was added because the enzyme of interest is suspected to be Zn^2+^ dependent. The purified protein was flash frozen using liquid nitrogen in 200 μl aliquots and subsequently stored at -80°C.

### β-Lactamase activity assay

The purified, soluble protein (MBX3358097.1) was tested with standard β-lactamase activity detection assays using the chromogenic cephalosporin analog, nitrocefin (Sigma Aldrich) [53]. Nitrocefin solutions were prepared in a 40 mM stock solution in DMSO and then diluted in 20 mM Tris buffer (pH = 7.0) immediately before use in a 1 mM working solution. Briefly, standard nitrocefin assays were performed in a 96-well plate by combining 180μl 20 mM Tris buffer with 0.1 mM ZnCl_2_ (pH = 7.0) with 10 μL purified enzyme and 10μL nitrocefin working solution. The plate was incubated in a plate reader (BioTek Synergy H1, Agilent) at 25°C, and the absorbance at 482 nm was measured every minute for 2 h. As negative controls, boiled enzyme and buffer-only conditions were included in every assay. To obtain the boiled enzyme control, aliquots of purified protein were incubated at 100°C for 15 minutes. Reaction rates were quantified from continuous measurements as described previously by Ryu et al. using the molar extinction coefficient of nitrocefin (12,124 M^−1^ cm^−1^) in 20 mM Tris [54].

## Results and discussion

### Biotic factors are more dominant than abiotic factors for altering the taxonomic and functional composition of biofilms grown in WWTP effluent

We deeply sequenced metagenomes from a total of 75 biofilm and water samples to an average sequencing depth of 138 million reads per sample (max. 204 million reads). After metagenomic assembly, we analyzed the taxonomic composition of stream biofilm communities to assess whether community composition was correlated with different experimental treatments: 30% WW, 80% WW, 30% UF WW, 80% UF WW, or stream water (0% WW). Experimental treatment was correlated with taxonomic composition for both prokaryotes (PERMANOVA, R^2^ = 0.65, p < 0.001) (**Figure 1**) and eukaryotes (PERMANOVA, R^2^ = 0.70, p < 0.001) (**Figures S1-S3**) at the genus level [30]. Growth in 80% WW relative to stream water (0% WW) altered prokaryotic biofilm community composition, namely we observed a decrease in the relative abundance of *Cyanobacteria* and *Verrucomicrobiota* and an increase in the relative abundance of *Bacteroidota* (**Figure 1**), in agreement with previous 16S rRNA gene-based analysis [27]. These shifts also mirror changes in microbial communities downstream of WWTPs relative to communities upstream [14]. However, these previous field studies did not include ultrafiltration treatment. We therefore compared WW UF and non-UF WW-grown bioiflms, and identified several bacterial phyla which showed significant shifts in relative abundance, namely a significant decrease in *Myxococcota* and an increase in *Planctomycetota* (**Figure 1**). UF treatment therefore altered microbial community composition including taxa with specific relevance for WWTP processes. For example, *Planctomycetota* is the only known phylum capable of anaerobic ammonia oxidation (anammox) for nitrogen removal from WW [25,55,56].

**Figure 1.**
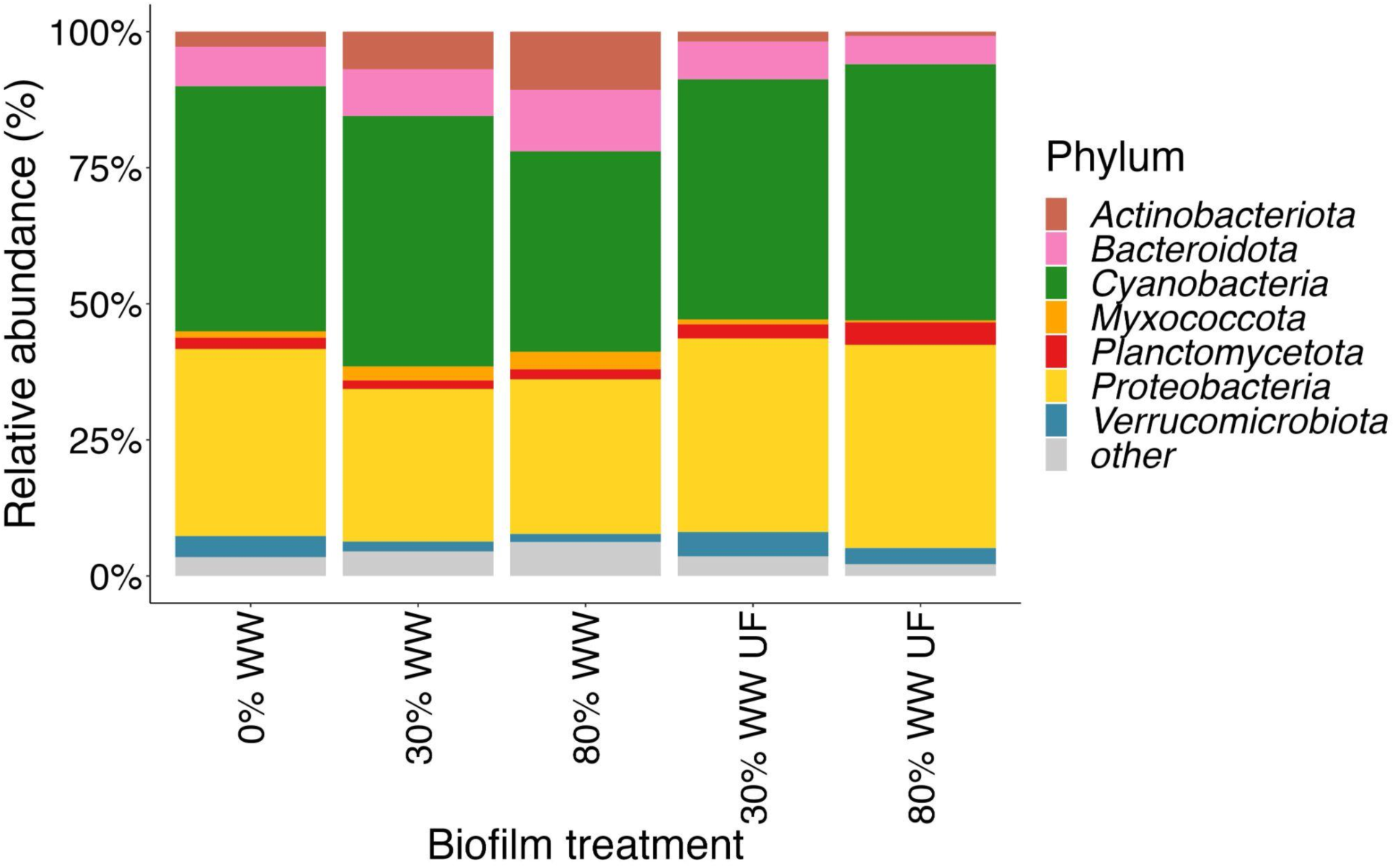
Mean relative abundance of prokaryotic phyla in the metagenomes of stream biofilms. Taxonomic shifts are shown for biofilms which were grown for four weeks continuously in stream water mixed with treated wastewater (WW) with or without ultrafiltration (UF) treatment. Taxa that were in the upper decile of abundance in at least one experimental treatment were retained while the remaining phyla were binned as ‘other’.

The taxonomic composition of biofilms grown in UF WW was more similar to that of biofilms grown in 0% WW than to biofilms grown in non-UF WW, suggesting that biotic factors from WWTPs play a greater role in changes in the stream biofilm community than abiotic factors (**Figure 2, Figure S3**). Metagenomic taxonomic analysis demonstrated that WW alters stream biofilm composition, supporting previous studies that some microorganisms from WW are able to colonize and modify biofilm community composition [27]. To leverage the shotgun metagenomic data, we investigated whether treatment also altered the composition of functional enzyme diversity (see Methods). PCoA revealed that the functional enzyme composition (**Figure S4**) was also significantly associated with the experimental treatment (PERMANOVA, R^2^ = 0.58, p < 0.001).

**Figure 2.**
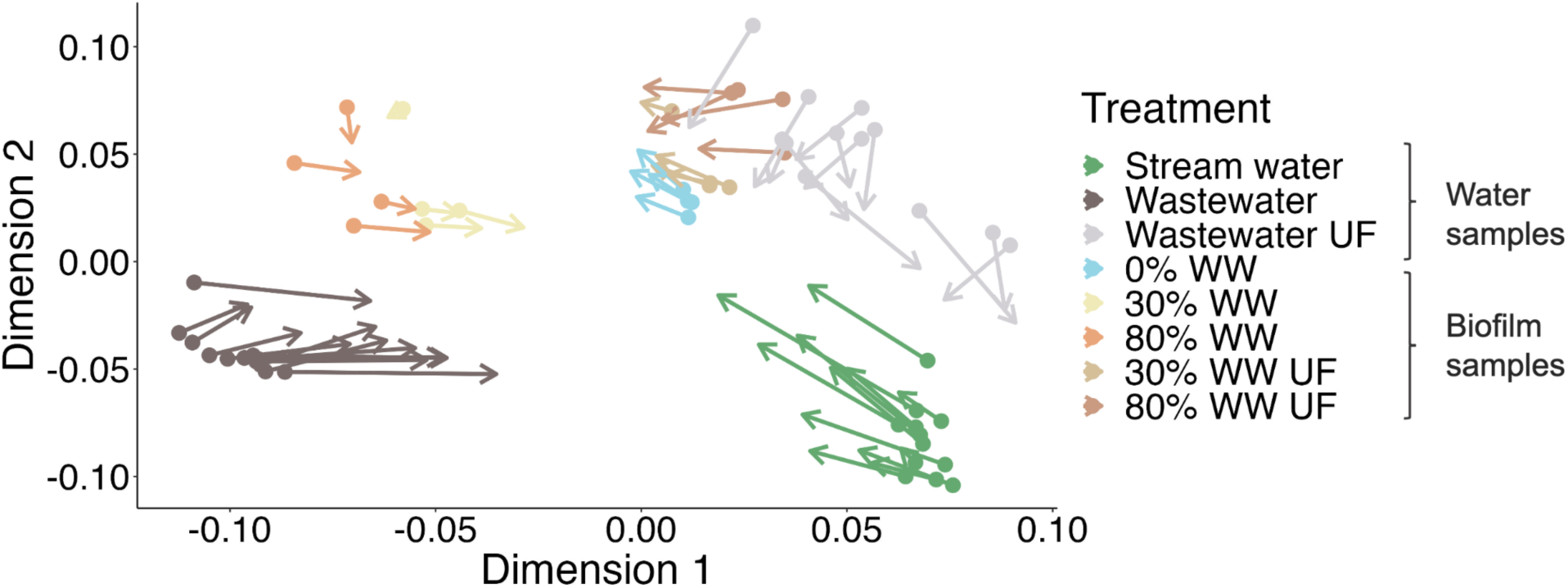
PCoA-based Procrustes analysis showing a structural alignment between microbiome and resistome composition. Points indicate positions on PCoA axes 1 and 2 of prokaryotic composition (microbiome), while arrow tips indicate positions on PCoA axes 1 and 2 of resistome composition following Procrustes rotation. The correlation coefficient in a symmetric Procrustes rotation was 0.901, the sum of squares (m12 squared) was 0.1889), and the p-value was less than 0.001.

### Resistome composition correlates with microbiome composition

We next examined how the total set of ARGs detected in each sample - the ‘resistome’ - varied with experimental treatment (e.g., WW% and non-UF vs. UF). Resistome composition was significantly affected by experimental treatment (PERMANOVA, R^2^ = 0.60, p < 0.001), similar to functional enzyme and taxonomic microbiome composition (**Figure S5**). Biofilms exposed to 30% or 80% UF WW clustered with 0% WW samples as opposed to samples exposed to equivalent amounts of non-UF WW (**Figure 2**).

To test for a relationship between the microbiome and resistome composition, we used Procrustes superimposition of resistome PCoA vectors onto prokaryotic microbiome PCoA vectors (**Figure 2**). Procrustes analysis revealed that the composition of the resistome was structurally related to the composition of the prokaryotic microbiome. The low Procrustes sum of squares of 0.1889 and calculated correlation value of 0.9 in a symmetric Procrustes rotation (p-value < 0.001) indicated a strong alignment between the resistome and microbiome. Similarly, functional enzyme composition was also associated with the resistome composition (**Figure S6,** Procrustes corr. coeff. = 0.89, p-value < 0.001). Alignments between the microbiome and resistome composition has been previously observed in microbial communities downstream of WWTPs [13,14], but local environmental features at different sampling sites are potential confounding factors that may have determined both microbiome and resistome composition in the field. Here, we demonstrated a microbiome-resistome relationship in a controlled experiment with biofilms grown in a defined gradient of stream water to WW and ultrafiltered WW matrices. Generally, the rotated PCoA vectors for the resistome are closer to the prokaryotic microbiome PCoA vectors for biofilm samples than they are for water samples (**Figure 2**). These results further suggest that changes in resistome composition in WW-impacted stream biofilms are likely due to changes in the biofilm communities by WWTP bacteria such as by colonization [27,57].

### WW ultrafiltration treatment has ARG class-specific effects on biofilm resistomes

A total of 570 ARGs across 41 different higher-level ARG classes were identified in the biofilm metagenomes using a short reads pipeline for ARG detection [34]. In biofilms, the abundances of ARGs conferring multidrug resistance, in addition to ARGs conferring resistance to beta-lactams, fluoroquinolones, peptides, rifamycins, and tetracyclines, differed significantly between conditions **(Table S5)**.

The normalized read counts of several, but not all, ARG classes increased with increasing WW%, as shown in **Figure 3**. In contrast, the metagenomic abundances of ARG classes did not increase in biofilms grown in comparable percentages of UF WW with the exception of sulfonamide AR Gs (**Figure 3**). These UF treatment results therefore suggest biotic factors primarily alter the abundances of ARGs in WW-grown biofilms.

**Figure 3.**
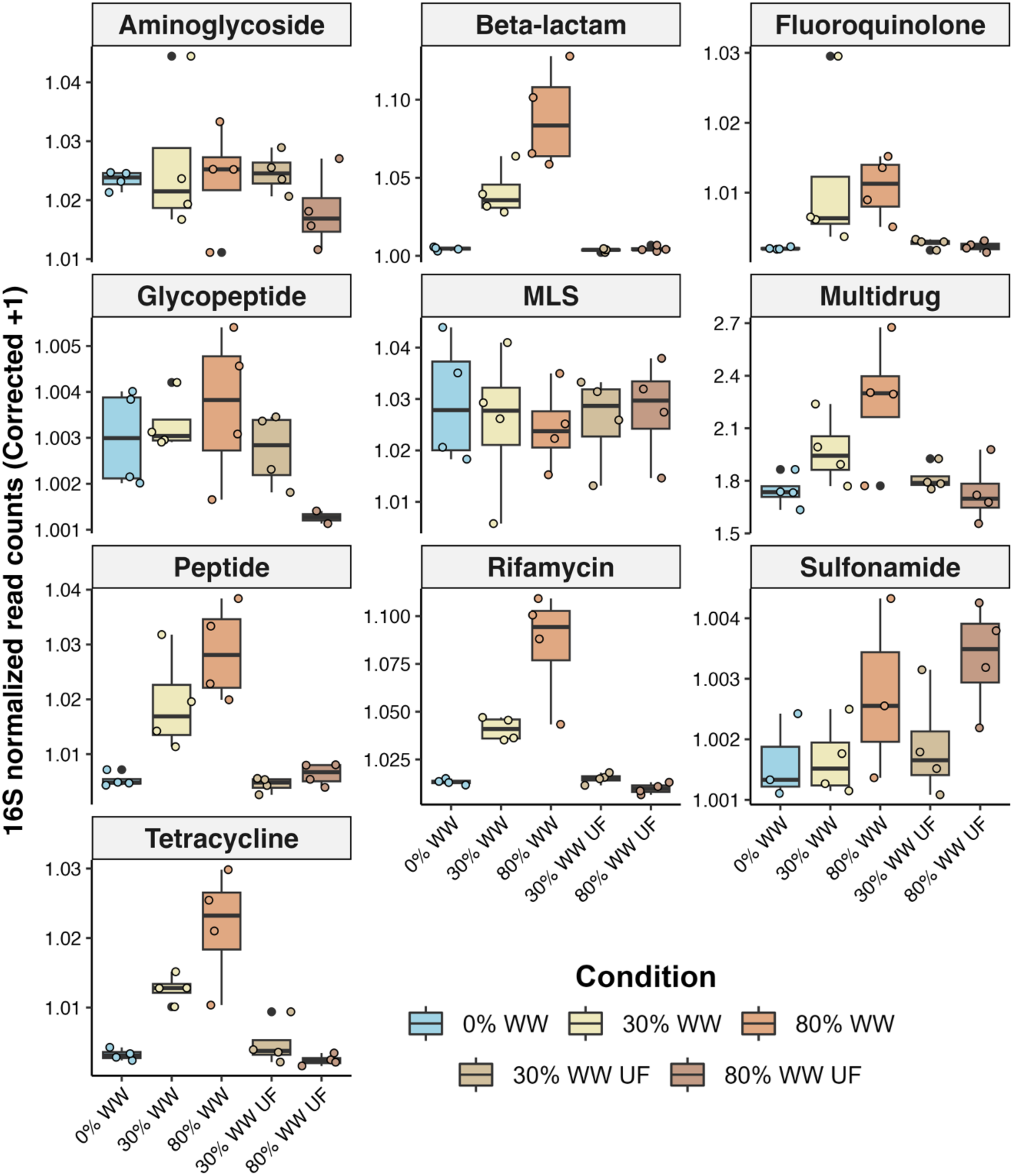
Significant ARG classes profiled using a metagenomic short reads pipeline. Abundance (16S-normalized read counts) of the DeepARG hits assigned to ARG classes that varied significantly with adjusted p-values ≤ 0.001 (Kruskal-Wallis test + Benjamini-Hochberg multiple hypothesis correction) between different treatment conditions. MLS = macrolide-lincosamide-streptogramin.

As a notable exception, sulfonamide ARGs exhibited increasing abundance with increasing WW% yet they displayed the highest abundance under 80% UF WW conditions, suggesting a contribution from abiotic factors such as nutrients or micropollutants (MPs) in WW. The high abundance of sulfonamide ARGs in UF-grown biofilms may also be related to gene family-specific mobility mechanisms such as extracellular genetic material which is small enough to pass through the UF membrane. In support of this hypothesis, the sulfonamide ARGs *sul1-4* are known to be highly mobilized in WWTP effluent microbiomes [58]. Although many ARGs are mobile, *sul1* ARGs are known for their pervasive colocalization within class 1 integrons [44] enabling bacteria to acquire and swap sulfonamide resistance genes [59,60]. Analysis of the assembled metagenomic contigs in our samples revealed that *sul1* genes were frequently colocalized on the same metagenomic contigs as mobile genetic elements, including *int1* integron integrases, as well as recombinases and transposases (**Figure S7**). Critically, the presence of *intI1* genes in environmental samples has been suggested as a gene proxy for estimating the levels of anthropogenic pollution, including MPs, as proposed by Gillings et al. [60]. The high abundance of sulfonamide ARGs observed in UF WW-grown biofilms supports the relationship between *int1* genes, sulfonamide ARGs typically associated with *int1,* and abiotic factors, including pollutants [59].

### Associations between assembled ARG abundances and WW percentages

Metagenomic assembly-based approaches are likely to have lower sensitivity but higher specificity than short read-based approaches in the detection of antibiotic resistance genes [61,62]. Therefore, as a complementary analysis to the DeepARG short read-based detection methods, we used stringent criteria (see Methods) to query our assembled metagenomes against the Structural Antibiotic Resistance Genes (SARG) database [43]. The query yielded 2,240 hits that clustered at the 70% amino acid sequence similarity level into 102 ARG ‘cluster representatives.’ Analysis by ARG class resulted in the following ARG cluster counts: 15 beta-

To quantify relative ARG abundances across treatment conditions, we mapped metagenomic reads back to the 102 ARG cluster representatives. Ten ARG cluster representatives differed significantly (Kruskal-Wallis test, p < 0.05) in abundance between at least one experimental treatment condition (**Table S6**). In addition to two beta-lactam resistance ARGs, one each of the multidrug, MLS, and aminoglycoside ARG representatives displayed the trend of increasing abundance with increasing WW%. The fold change increase in metagenomic abundance was the greatest for two beta-lactam resistance ARG clusters (Class-A beta-lactamase gene, p = 0.0054; and a Class-D beta-lactamase bla OXA-4 gene, p = 0.0054; **Figure 4**). In contrast, a third beta-lactam ARG (extended-spectrum BEL family beta-lactamase, p-value = 0.0488) displayed high coverage exclusively in the 80% UF WW condition (**Figure 4**). These results suggest that the abundance of this cluster of BEL family ARGs may be influenced by abiotic factors such as MPs, the effects of which may have been outcompeted by biotic effects in the non-UF WW-grown biofilms. Leveraging our metagenomic assembly-based approach, we examined genes localized on the same metagenomic contig as the BEL family beta-lactamase hit. The BEL family beta-lactamase is directly flanked by a *Pseudomonas* TniC resolvase. This TniC resolvase shares 100% sequence identity with a TniC resolvase (ADC80450.1) in integron/gene cassette systems previously shown to be influenced by abiotic factors (heavy metal concentrations [63]. Although the causality must be demonstrated, genetic mobility influenced by abiotic factors may contribute to the observed abundance of BEL family beta-lactamases in 80% UF WW-grown biofilms.

**Figure 4.**
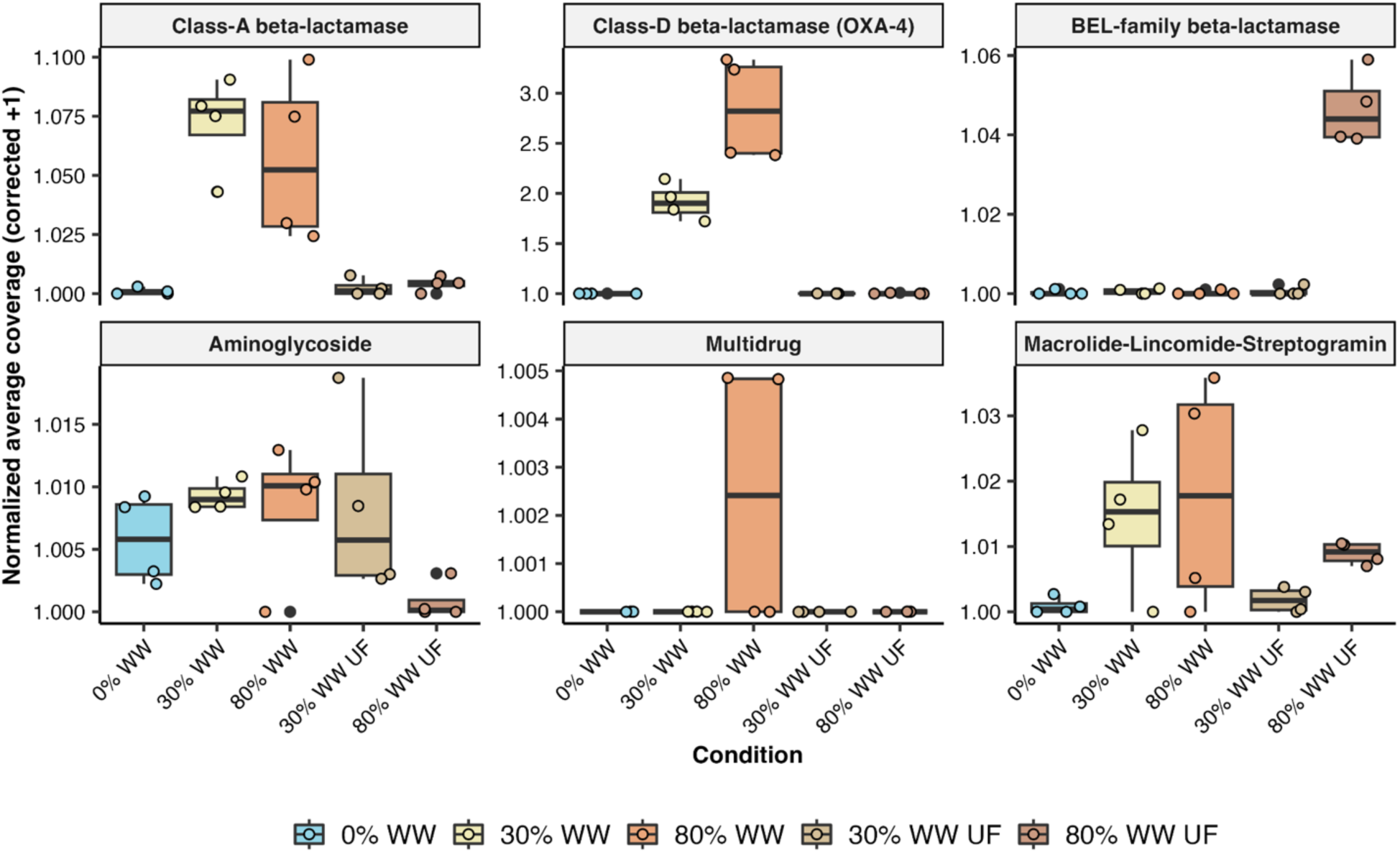
Significant ARG classes using a metagenomic assembly-based approach. The corrected normalized average coverage of ARG cluster representatives identified using an assembly-based ARG detection method. ARG clusters with significant differences in abundance between at least one different treatment condition are displayed (adjusted p-values ≤ 0.05 using Kruskal-Wallis test + Benjamini-Hochberg multiple hypothesis correction).

Overall, our experimental design separated biotic and abiotic factors and shed light on the gene family-specific mechanisms underlying how WWTP effluent may alter the resistome composition of stream biofilms. Our combined results from short read mapping and assembly-based ARG analysis generally aligned with the exception of the sulfonamide ARGs, which did not show a significant difference in the assembly-based approach. This may be due to limitations in the metagenomic assembly and highlights the value of using two different, complementary approaches. For example, the significance of beta-lactamase ARGs identified via the short read-based approach was also captured via the assembly-based approach. Notably, a consensus for beta-lactamase abundance emerged from the comparison of our study to previous studies in Switzerland [13] and globally [9,14,64,65]. Beta-lactamase ARGs consistently remain the most abundant class of ARGs detected in WWTP effluent and aquatic environments [7,66–69]. Their prevalence can be attributed to a variety of factors, including the early discovery and continued widespread overuse of beta-lactam antibiotics for humans and animals [70,71]. However, the unexpectedly high abundance of BEL family beta-lactamases exclusively in 80% UF WW samples (**Figure 4)** prompted us to focus on beta-lactamases for further experimental investigation and analysis of abiotic factors.

### Micropollutants as abiotic factors and their potential influence on ARG transfer

We examined MP concentrations as abiotic factors capable of influencing gene family-specific ARG abundances, such as the *sul1-4* and BEL family beta-lactamase ARGs showing unexpectedly higher abundances in the UF WW compared to non-UF conditions (**Figure 3**, **Figure 4)**. Based on previous chemical analysis by liquid chromatography-tandem mass spectrometry [11], the MP concentrations measured in the flume biofilm samples increased with increasing WW% (**Table S4**), but the concentrations of MPs extracted from biofilms did not significantly differ between the UF and non-UF WW. Many of the antibiotics measured, including sulfamethoxazole and sulfapyridine (**Table S4**), were present at concentrations less than or equal to the limit of quantification in WW-grown biofilms. Among all antibiotics detected, the macrolide clarithromycin exhibited the highest concentrations in 80% WW and 80% UF WW grown biofilms (45-95 ng per mg biofilm, **Table S4**). In line with the general MP trends, clarithromycin concentrations increased with WW% and did not vary significantly between UF and non-UF treatments. In contrast, the metagenomic abundances of macrolide-lincosamide-streptogramin (MLS) ARGs, which typically confer resistance to clarithromycin, were significantly lower in UF than in non-UF conditions (**Figure 4**). This represents a case of an antibiotic-ARG pair where ARG abundances did not correlate with the expected abiotic factor (clarithromycin). However, this does not take into account the potential effects of other MPs which are not classified as antibiotics but nonetheless influence ARG abundance [3].

A growing body of evidence indicates that many anthropogenic pollutants that do not have known antimicrobial bioactivities nonetheless influence ARG transfer and proliferation [3,72–77]. However, there is generally a lack of knowledge on the mechanisms underlying how MPs without antibiotic activities act as selective agents for ARG abundance. In our study, the 75 different MPs measured [11] included artificial sweeteners, non-antibiotic pharmaceuticals, corrosion inhibitors, and pesticides (**Table S4)**. Compared with the other compound classes, artificial sweeteners were the most abundant MPs, with concentrations as high as µg/L detected in WW samples [11] (**Table S4**). Notably, the artificial sweeteners measured in the biofilms (acesulfame K, cyclamate, and saccharin) all contain sulfonamide moieties which are chemically analogous to the moieties found in sulfonamide antibiotics (**Figure S8**). Sulfonamides were among the few ARG classes with significantly greater abundances in UF WW-grown biofilms than in non-UF WW-grown biofilms (**Figure 3**). These observations led us to speculate that abiotic factors, such as sulfonamide-containing MPs, may play a role in promoting the abundance of such gene family-specific ARGs. In support of this, several experimental studies have shown that artificial sweeteners promote high rates of horizontal gene transfer and natural uptake of ARGs in bacteria [76,78]. Further work has demonstrated that acesulfame-K and other artificial sweeteners directly enhance the rate and quantity of horizontal plasmid-mediated transfers of *sul* genes [79]. Interestingly, a plasmid-encoded microbial hydrolase capable of cleaving the sulfonamide moiety of acesulfame-K was recently identified and shown to have a metallo beta-lactamase structural fold [80]. Taken together, these findings prompted us to conduct a deeper functional investigation of beta-lactamase fold genes whose abundance increased with WW%.

### *Planctomycetota* beta-lactamases display differential abundances in response to sulfonamide-containing MPs and WWTP effluent

To investigate the relationship between ARG abundances and sulfonamide-containing MPs, we consulted previous reports of enrichment studies of activated sludge communities grown on sulfonamide-containing MPs including artificial sweeteners [81,82]. Multiple independent enrichment studies on acesulfame-K [82], saccharin [81], and cyclamate [81] reported significant increases in the relative abundance of *Planctomycetota* and particularly the family *Phycisphaeraceae*. Previously, Desiante et al. (2022) also identified significant differences in the relative abundances of *Planctomycetota* and *Phycisphaeraceae* in biofilms grown in 80% WW and 80% UF WW in two independent flume biofilm experiments (log_2_ -fold change > 4; p < 0.05). We therefore conducted differential abundance analysis and observed statistically significant differences in the *Planctomycetota* relative abundances between WW and UF WW treatments (**Figure 5A**). Taxa assigned to the phylum *Planctomycetota* constitute up to 2.1% of the relative abundance of our biofilm, WW, and stream water metagenomes, confirming previous reports of the *Planctomycetota* as a numerically-important constituent both freshwater and wastewater microbial communities [25,26,56]. Many *Phycisphaeraceae* are algae-associated with the metabolic potential to impact the fitness of their algal hosts [83], but the current unculturability of most planctomycetes limits our knowledge of linking their ARG genotypes to their phenotypes. To address this knowledge gap, we chose to target *Phycisphaeraceae* ARGs for experimental characterization.

**Figure 5.**
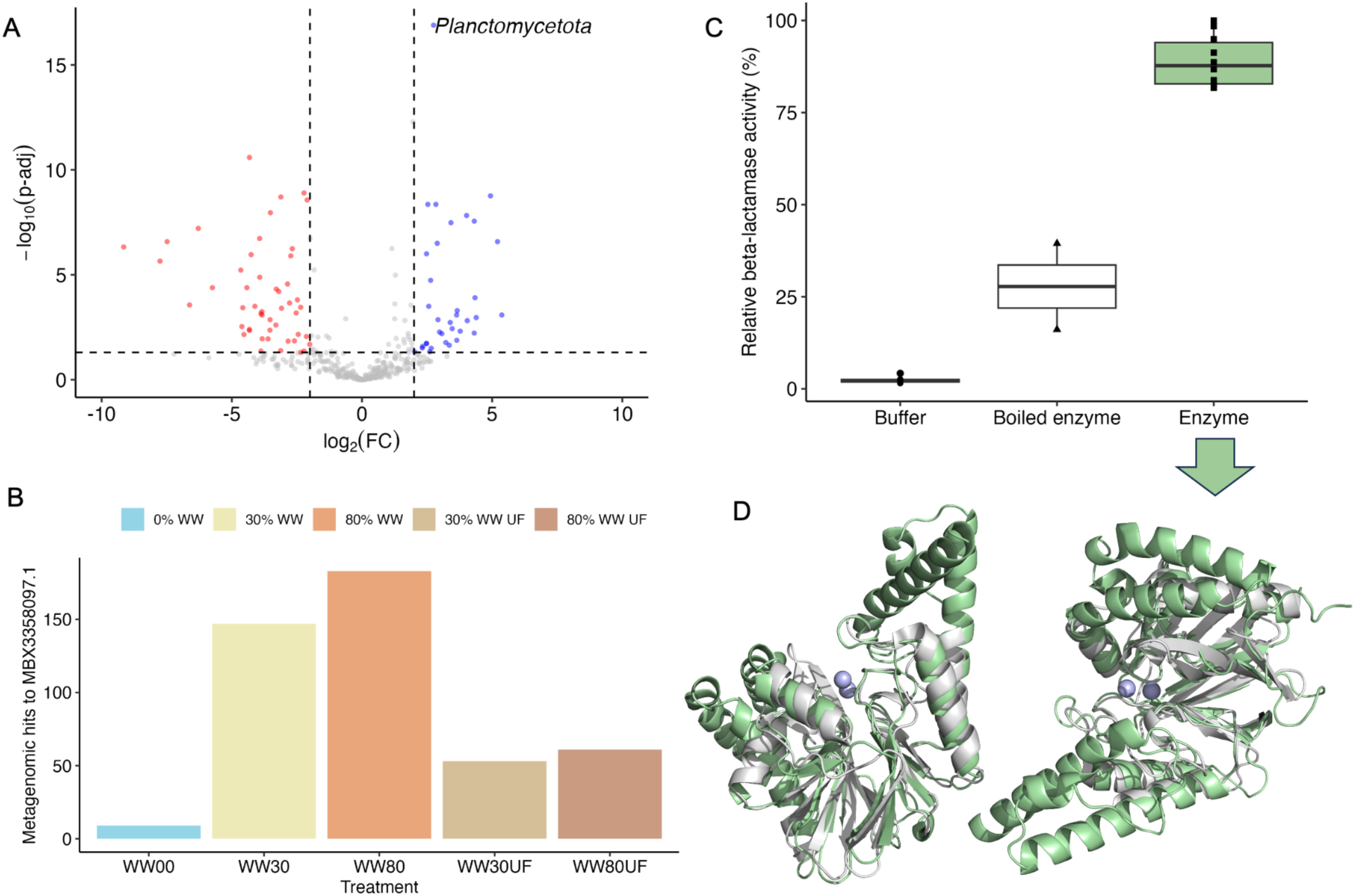
Functional characterization of an active *Planctomycetota* beta-lactamase. A) Volcano plot displaying the log_2_-fold change of differentially abundant taxa between WW30UF vs. WW30. B) Counts of hits showing significant homology to the putative beta-lactamase MBX3358097.1 in biofilm metagenomes. C) Purified enzyme assays demonstrated beta-lactamase activity of MBX3358097.1 with the colorimetric cephalosporin analog, nitrocefin, compared to boiled enzyme and buffer only controls. D) Two views of a high-confidence AlphaFold2 model (mean pLDDT = 96.5) of MBX3358097.1 structurally-aligned with its closest structural homolog in the Protein Data Bank (PDB100), as identified by FoldSeek [85] as of November 2023, the *Bacillus cereus* beta-lactamase (PDB ID: 3KNS).

Due to known challenges in planctomycete cultivation [56], we used heterologous expression techniques to functionally validate two *Phycisphaeraceae* ARGs. We focused specifically on *Phycisphaeraceae* beta-lactamases for four reasons: (1) the beta-lactamases were identified in our ARG analysis as the most abundant and significantly-different class of ARGs affected by the percentage of treated wastewater, (2) beta-lactamase fold enzymes were also previously identified as plasmid-encoded enzymes active in the biotransformation of the sulfonamide MP acesulfame-K [80], revealing the ‘structural overlap’ of a versatile metallo beta-lactamase fold capable of cleaving both sulfonamide- and beta-lactam moieties, and (3) previous evidence of sulfonamide cleavage in the sweetener cyclamate was shown by enzymes which were metal-dependent hydrolases but their sequences were not determined [84]. Finally, (4) full-length homologs with greater than 50% amino acid identity to the sulfonamide resistance genes *sul1-4* were not detected in the *Phycisphaeraceae*. Since classical sulfonamide resistance genes were lacking, we hypothesized that metallo beta-lactamase-fold enzymes may instead play an overlooked role in sulfonamide cleavage in the *Phycisphaeraceae*.

### Functional characterization of a new beta-lactamase ARG from the *Planctomycetota*

To identify metallo-beta-lactamase fold enzymes assigned to the *Phycisphaeraceae*, we queried the 75 metagenomes sequenced in this study as well as additional publicly-available metagenomes from previous sulfonamide-containing MP enrichment studies [81,82]. Through bioinformatic prioritization (Methods), we selected two full-length metallo beta-lactamase fold protein hits from sulfonamide-enriched *Phycisphaeraceae* MAGs for functional characterization [82]. The abundances of both proteins increased with increasing WW% (**Figure 5B** and **Figure S9**). We heterologously expressed each of the *Phycisphaeraceae* genes in *E. coli.* One protein did not express in *E. coli* and despite troubleshooting was not amenable to protein production or purification with nickel-affinity chromatography. The second protein (MBX3358097.1) was soluble and purified to homogeneity with an average yield of 500 µg/mL protein (**Figure S10**) therefore we proceeded with the functional analysis of this protein. We tested the purified *Phycisphaeraceae* protein for beta-lactamase activity and observed enzyme-mediated cleavage of the cephalosporin analog, nitrocefin relative to boiled enzyme controls (**Figure 5C**). We next tested its substrate specificity for sulfonamide-containing compounds. By high-performance liquid chromatography analysis and comparison to authentic standards, we determined that the *Phycisphaeraceae* enzyme did not biotransform the sulfonamide-containing MPs acesulfame-K, saccharin, or cyclamate, nor did it transform sulfamethoxazole, thus disproving our hypothesis for its role in sulfonamide cleavage. Nonetheless, this analysis resulted in the first biochemical characterization of a new, active beta-lactamase from the *Phycisphaeraceae*.

MBX3358097.1 displayed a maximum 30% amino acid identity and between 20-50% query coverage with proteins of known function in the UniProtKB-SwissProt [86] and the Beta-Lactamase database (BLDB) [87]. These low sequence similarities limit reliable functional annotation and beta-lactamase classification based on homology alone. Therefore, we generated an AlphaFold2 model (Figure 5D, model available on GitHub) for the structural alignment and identification of beta-lactamase subfamily specific motifs [88,89]. MBX3358097.1 encodes a zinc-binding motif 2 (HxHxD**H**), which has a terminal histidine characteristic of the subclass B3 metallo beta-lactamases. However, in another known motif 4, this enzyme contains the residues ‘GD’, which are more characteristic of related glyoxalase II enzymes rather than B1-B3 family beta-lactamases [88,89]. These results are suggestive of promiscuous activity or additional enzymatic functions of MBX3358097.1 to investigate which are beyond the scope of this current study. Nonetheless, this analysis highlights the benefits for structural modeling of metagenomic sequences [90,91], particularly in cases where low sequence identity prevents reliable functional annotation

There are few previous reports of ARGs from the phylum *Planctomycetota* [92]. Planctomycetes were previously reported to have an unusual proteinaceous cell wall composition devoid of peptidoglycan [93], however, later findings demonstrated that planctomycetes do indeed produce peptidoglycan and are thus susceptible to cell-wall targeting antibiotics including beta-lactams [94]. In fact, a study of six culturable planctomycetes revealed that all of the strains tested were resistant to beta-lactam antibiotics although beta-lactamases could not be detected [95]. Importantly, MBX3358097.1 was also not detected by standard computational methods for detecting ARGs, including queries to the SARG [43] and CARD [96] databases due to its low sequence identity with known ARGs. This work therefore highlights the size of the knowledge gap and invites further inquiry into the role of sequence-diverse ARGs from uncultivated microorganisms and their contributions to environmental ARG reservoirs.

## Conclusions

Our results from a controlled flume system showed that WWTP effluent altered both the microbiome and resistome composition of stream biofilms. Through ultrafiltration treatment of WWTP effluent, we could attribute changes in ARG abundances primarily to biotic rather than abiotic factors. However, the abundance of several gene family-specific ARGs, most notably sulfonamides and beta-lactamases, suggested an influence by abiotic factors. We biochemically characterized the beta-lactamase activity of a new *Phycisphaeraceae* protein with low sequence identity to known ARGs. This provided proof-of-principle of a functional ARG from an uncultivated microorganism that was overlooked by standard computational ARG detection methods. This further highlights the need to characterize new ARGs from understudied taxa and to create a more comprehensive ARG catalog for querying complex microbiomes. Overall, our study exemplified a workflow to disentangle antibiotic resistance determinants in aquatic environments.

## List of abbreviations

ARGs: Antibiotic resistance genes
DNA: Deoxyribonucleic acid
MPs: Micropollutants
ORFs: Open reading frames
PCoA: Principal coordinate analysis
PCR: Polymerase chain reaction
rRNA: ribosomal Ribonucleic acid
SARG: Structured antibiotic resistance genes database
UF: Ultrafiltration
WW: Wastewater effluent
WWTPs: Wastewater treatment plants

## Ethical approval and consent to participate

Not applicable

## Availability of data and material

Metagenomic sequencing data generated during the current study have been deposited in the European Nucleotide Archive with the study accession code: PRJNA1008123. Scripts and additional data used in analysis and figures are available at:: https://github.com/MSM-group/AMR-ecoimpact-paper.

## Author’s contributions

M.A. contributed to the conceptualization, methodology, investigation, writing—editing and revisions of the manuscript, visualization, and funding acquisition. M.S. contributed to conceptualization, methodology, visualization, investigation, data curation, writing—editing and revisions of the manuscript. M.E. contributed to the methodology, investigation, and revision and editing of the manuscript. G.G. contributed to methodology and editing and revision of the manuscript. H.B. contributed to the conceptualization, and editing and revision of the manuscript. P.N.L.L. contributed to the revision and editing of the manuscript and project administration. K.F. contributed to the conceptualization, revision and editing of the manuscript and funding acquisition. J.v.d.V. contributed to the methodology, writing-editing and revision of the manuscript and project administration. S.L.R contributed to conceptualization, methodology, Investigation, writing-editing and revision of the manuscript, visualization, project administration and funding acquisition.

## Acknowledgements

We thank Dr. Bogdan Iorga for his export support in beta-lactamase classification and bioinformatic analysis. Robert Niederdorfer is acknowledged for his support and expertise in the metagenomic assembly through computational resources and collaboration with the Genetic Diversity Centre (GDC), ETH Zurich. We acknowledge Niklas Ferenc Trottmann for his training and support for the cloning, expression, and protein purification. We thank the Eawag Partnership Program together with the Institute of Water Education (IHE Delft) under the auspices of UNESCO for supporting the exchange of Mustafa Attrah to Eawag. We thank Elia Ceppi for assistance in interpretation of chemical micropollutant data and Thierry Marti for support in biological interpretation. We further acknowledge Stuart Dennis for his computational infrastructure support at Eawag. We acknowledge the EcoImpact teams 1.0 and 2.0 for fruitful discussions as well as conceptualizing and running the flume experiments which provided the samples necessary for this study.

