## Supplementary Information for "Disentangling abiotic and biotic effects of treated wastewater on stream biofilm resistomes enables the discovery of a new planctomycete beta-lactamase"

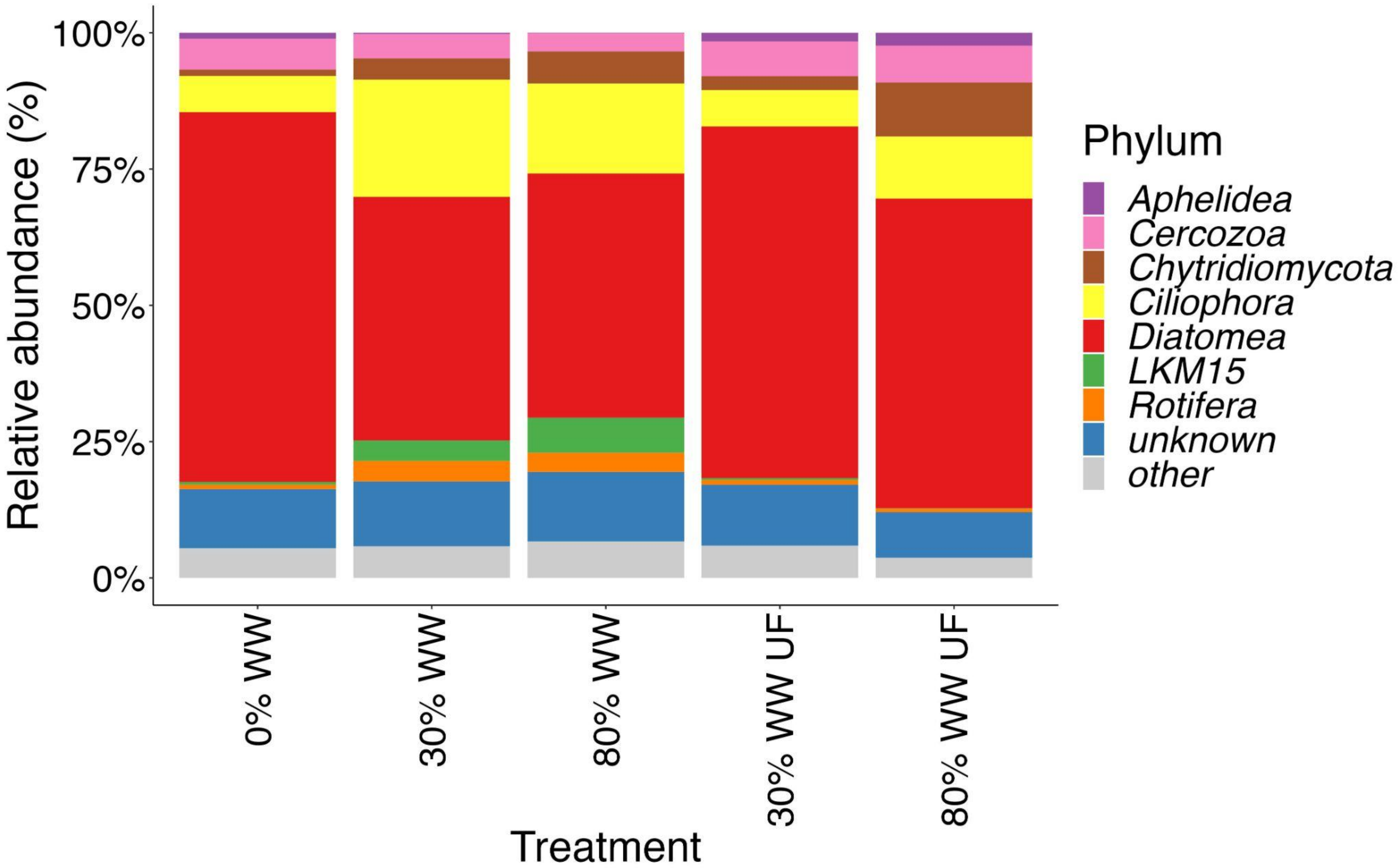


Figure S1. Mean relative abundance of eukaryotic phyla in each experimental treatment. Taxa which were in the upper decile of abundance in at least one experimental treatment were retained, while remaining phyla were binned as other.

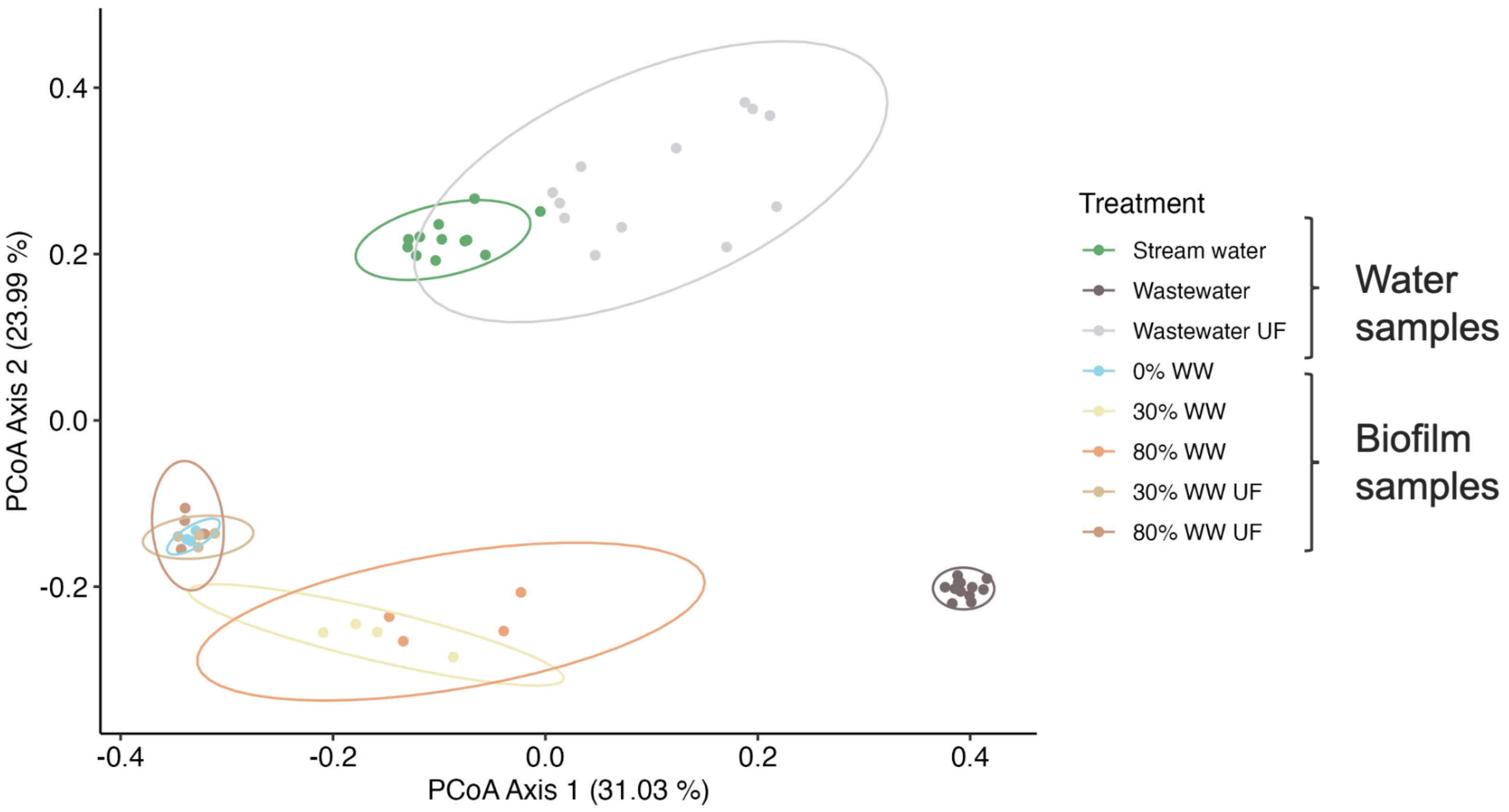


Figure S2. PCoA of Bray-Curtis distances between samples based on eukaryotic genus level abundance table inferred from the 75 metagenomes using mTags [30], with samples colored according to treatment. Samples with % WW or % WW UF indicated are biofilm samples, while remaining samples are water samples (95% confidence ellipses for different treatments).

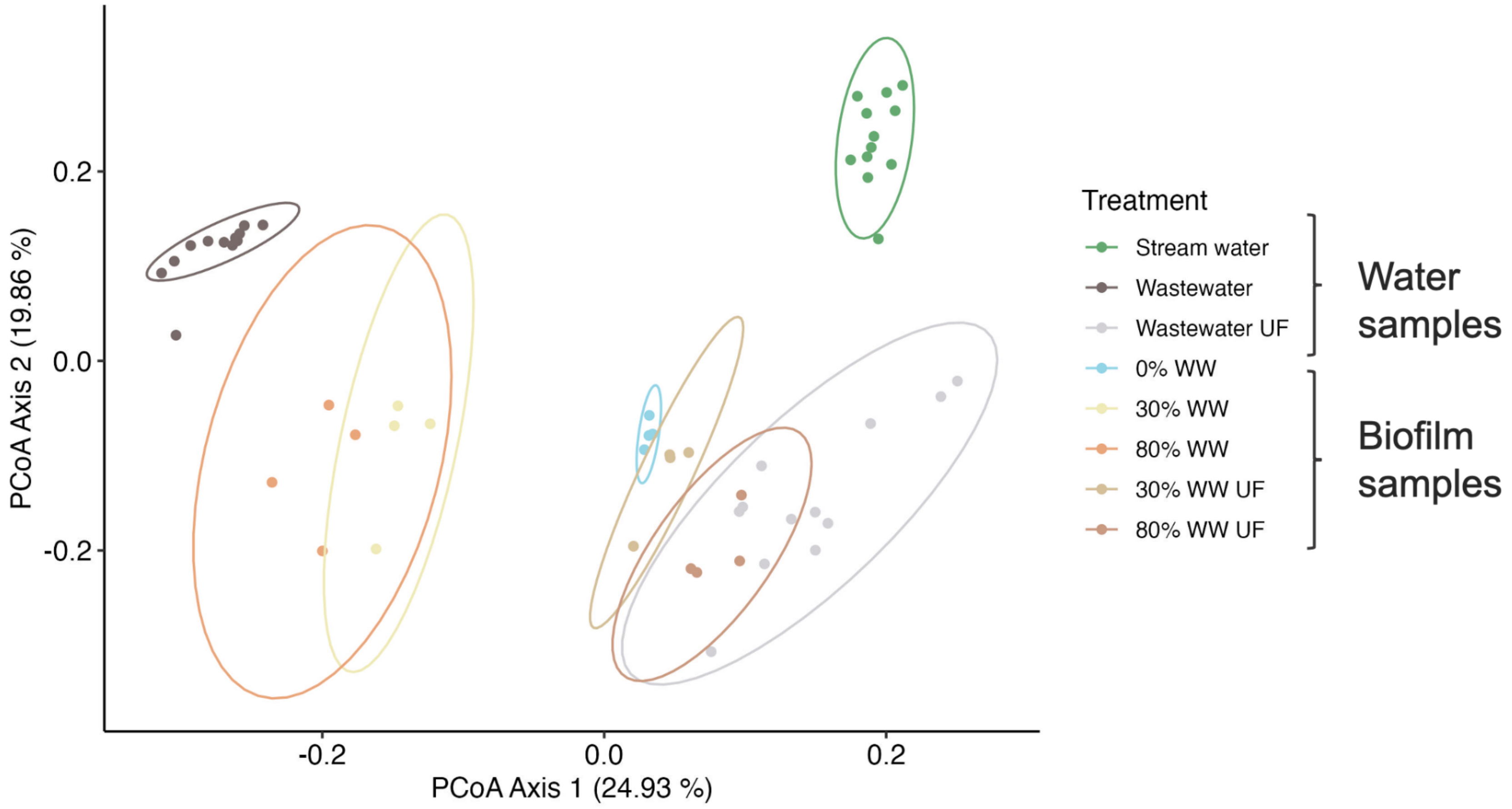


Figure S3. PCoA of Bray-Curtis distances between samples based on prokaryotic genus level abundance tables inferred from the 75 metagenomes using mTAGs [30], with samples colored according to treatment. Samples with % WW or % WW UF indicated are biofilm samples, while remaining samples are water samples (95% confidence ellipses for different treatments).

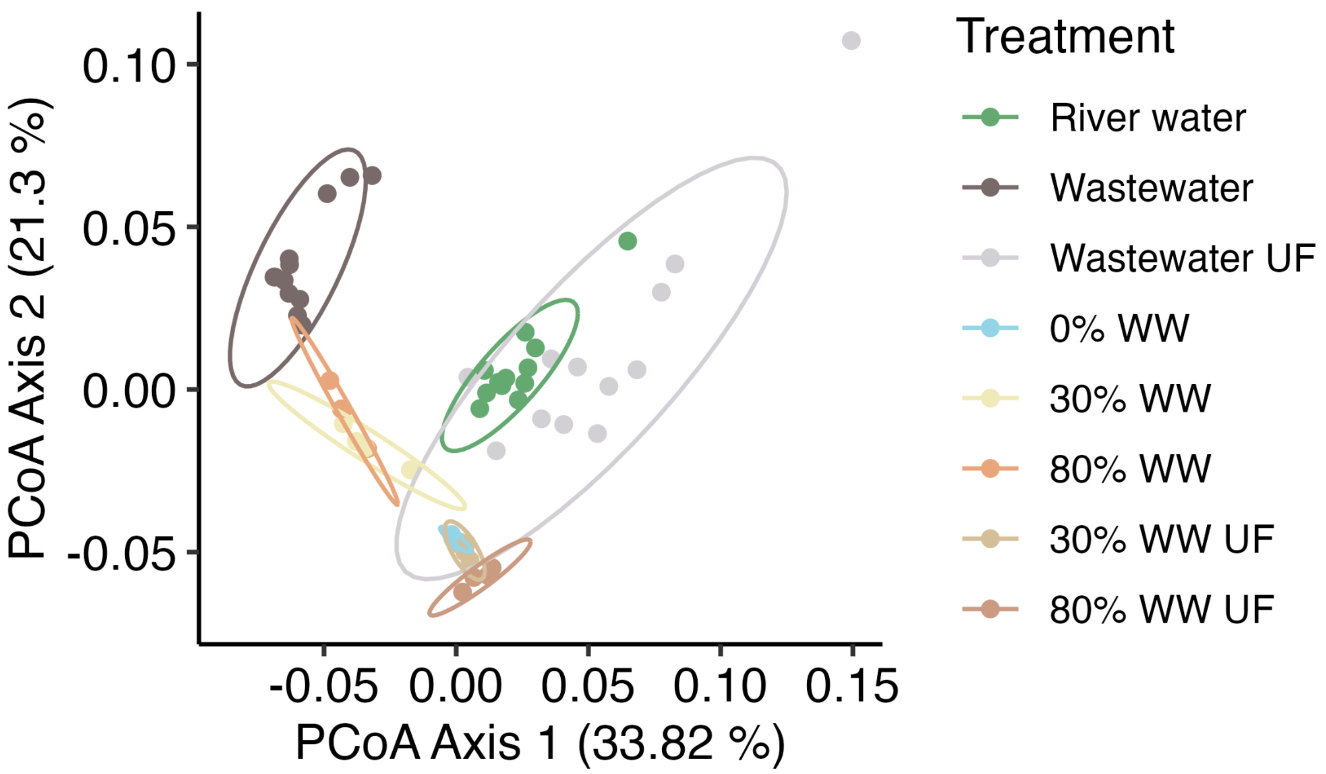


Figure S4. Principal coordinate analysis (PCoA) of functional enzyme counts across treatment conditions based on a Hellinger-transformation of counts of enzyme classes annotated with Enzyme Commission (EC) number fourth-level annotations (see Methods). The Bray-Curtis distance matrix was used (95% confidence ellipses for different treatments). Samples with % WW or % WW UF indicated are biofilm samples, while remaining samples are water samples (95% confidence ellipses for different treatments).

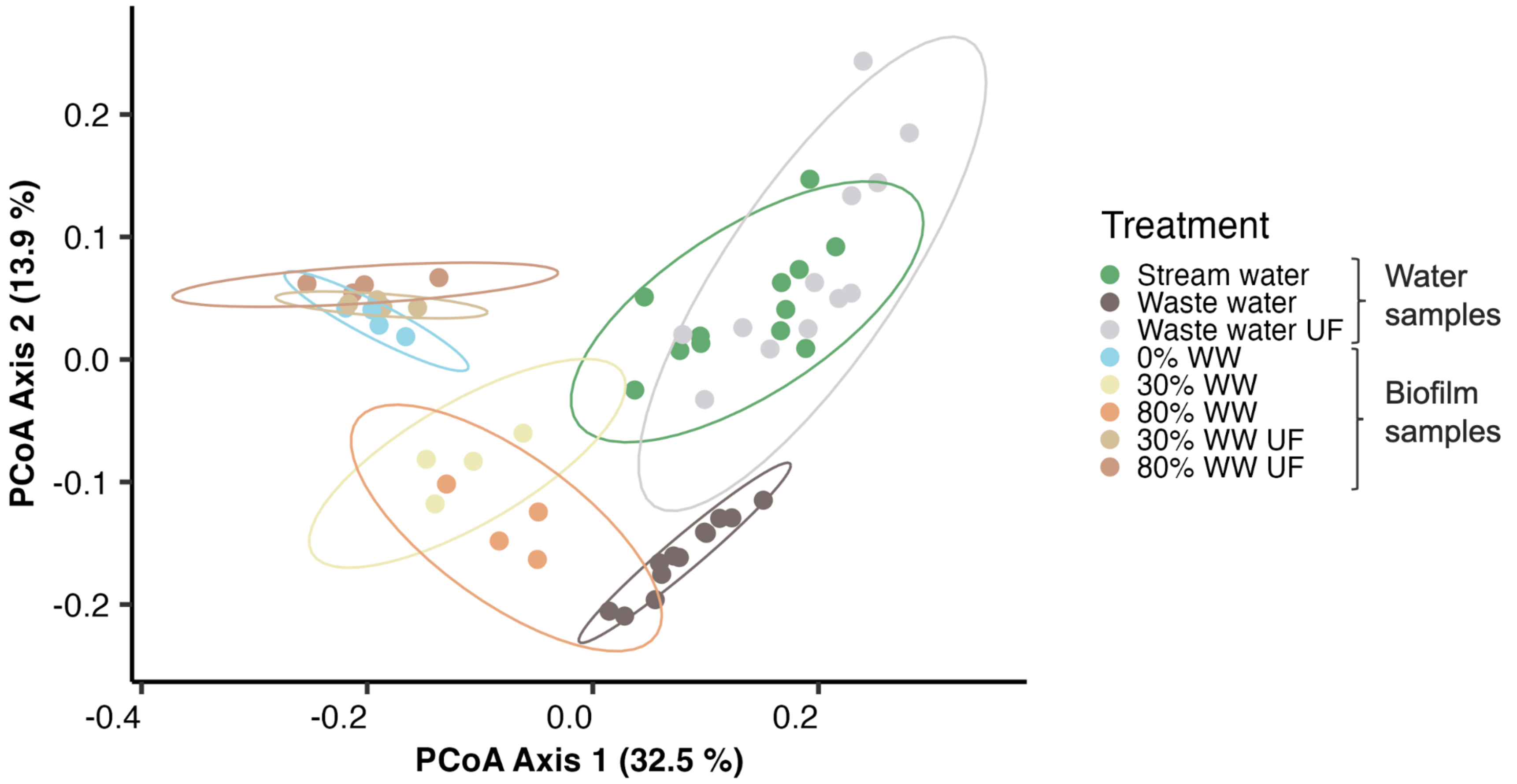


Figure S5**.** Principal coordinate analysis (PCoA) of DeepARG subtype counts across treatment conditions. ARGs were normalized by 16S read counts and the Bray-Curtis distance matrix was used (95% confidence ellipses for different treatments).

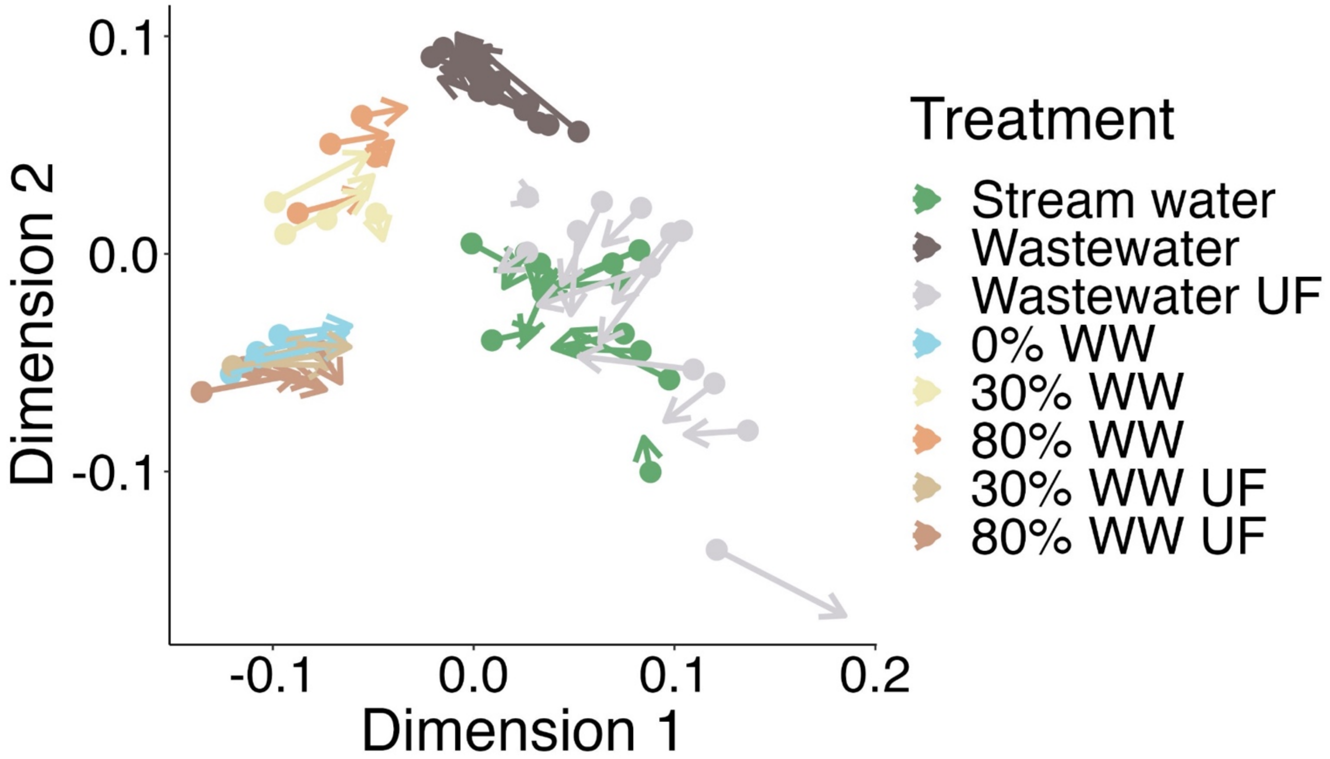


Figure S6. PCoA-based Procrustes analysis showing a structural alignment between resistome composition and functional enzyme composition (EC number, fourth-level) and antibiotic resistance profiles. Points indicate position on PCoA axes 1 and 2 of resistome composition, while arrow tips indicate position on PCoA axes 1 and 2 of functional genes corresponding to fourth-level EC counts were subjected to a Hellinger transformation followed by a Procrustes rotation. The correlation coefficient in a symmetric Procrustes rotation was 0.8901, the sum of squares (m12 squared) was 0.2078, and p-value was less than 0.001.

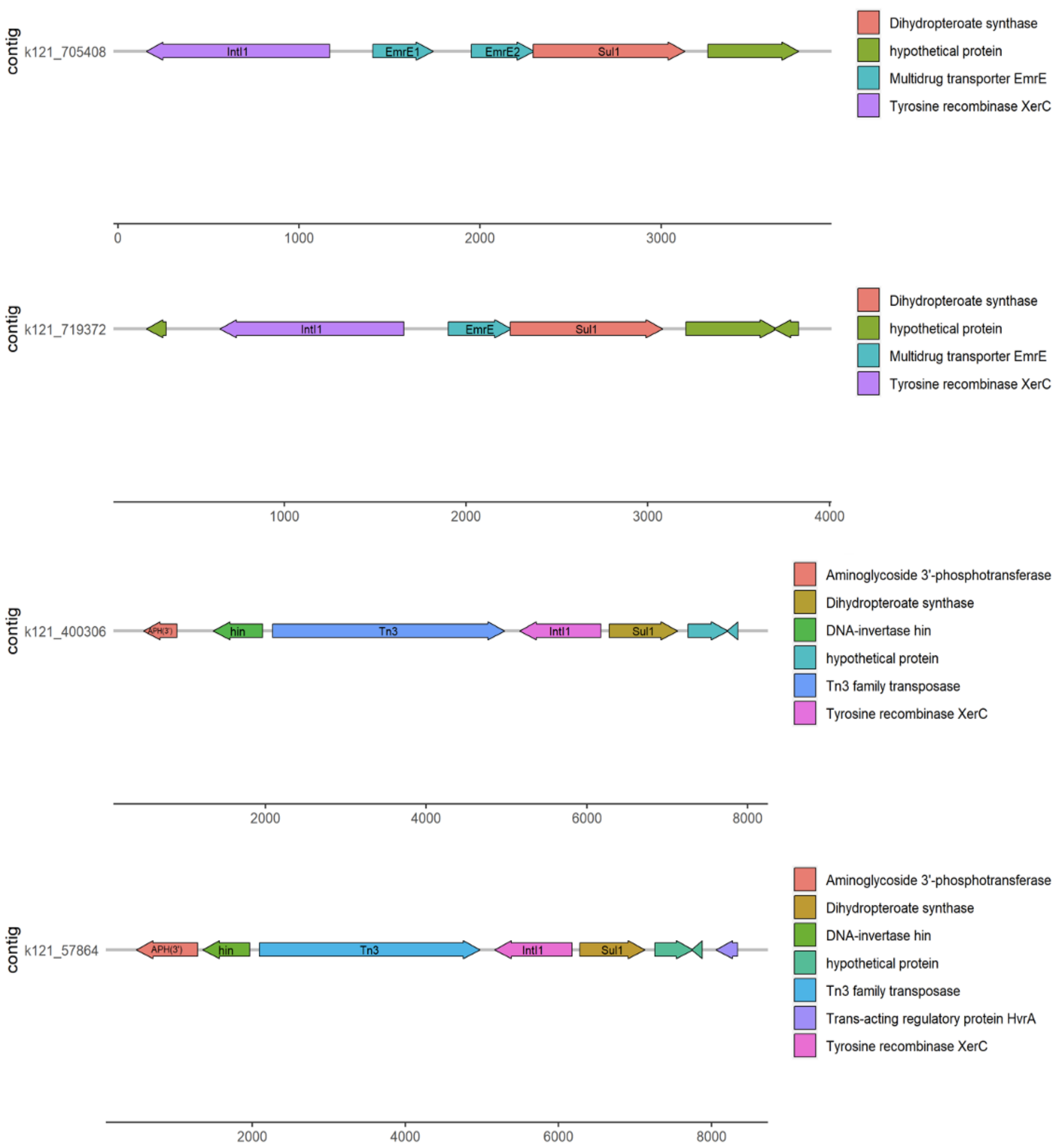
contig
k121_705408
IntI1
EmrE1
EmrE2
Sul1
0
1000
2000
3000
Dihydropteroate synthase
hypothetical protein
Multidrug transporter EmrE
Tyrosine recombinase XerC
contig
k121_719372
IntI1
EmrE
Sul1
1000
2000
3000
4000
Dihydropteroate synthase
hypothetical protein
Multidrug transporter EmrE
Tyrosine recombinase XerC
contig
k121_400306
APH(3')
hin
Tn3
IntI1
Sul1
2000
4000
6000
8000
Aminoglycoside 3'-phosphotransferase
Dihydropteroate synthase
DNA-invertase hin
hypothetical protein
Tn3 family transposase
Tyrosine recombinase XerC
contig
k121_57864
APH(3')
hin
Tn3
IntI1
Sul1
2000
4000
6000
8000
Aminoglycoside 3'-phosphotransferase
Dihydropteroate synthase
DNA-invertase hin
hypothetical protein
Tn3 family transposase
Trans-acting regulatory protein HvrA
Tyrosine recombinase XerC

Figure S7. Representative examples of *sul1* genes detected on biofilm metagenomic contigs co-localized with mobile genetic elements such as recombinases, integron-integrase, and transposases.

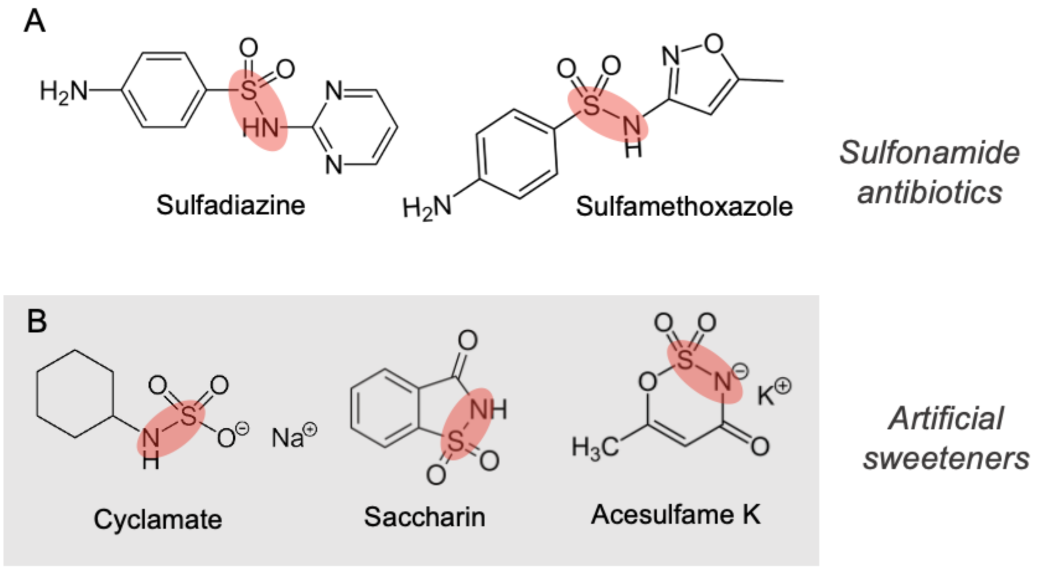


Figure S8. Chemical similarity between example A) sulfonamide-containing antibiotics and other B) sulfonamide-containing micropollutants detected in WW.

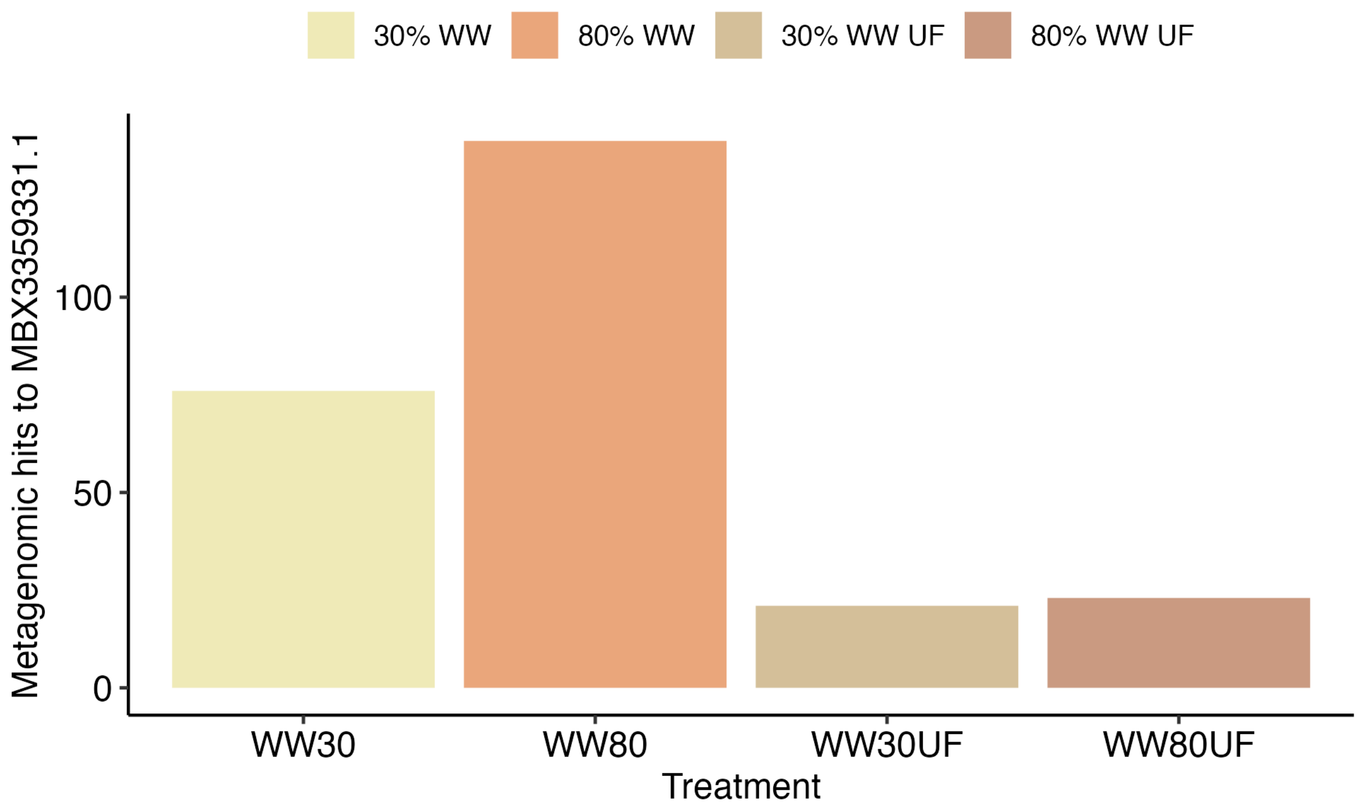


Figure S9. Metagenomic hits to the *Phycisphaeraceae* metallo beta-lactamase fold protein MBX3359331.1 shows higher counts in WW80 rather than WW30 and lower counts in the UF treated samples. No hits were obtained in the WW00 (stream water) samples.

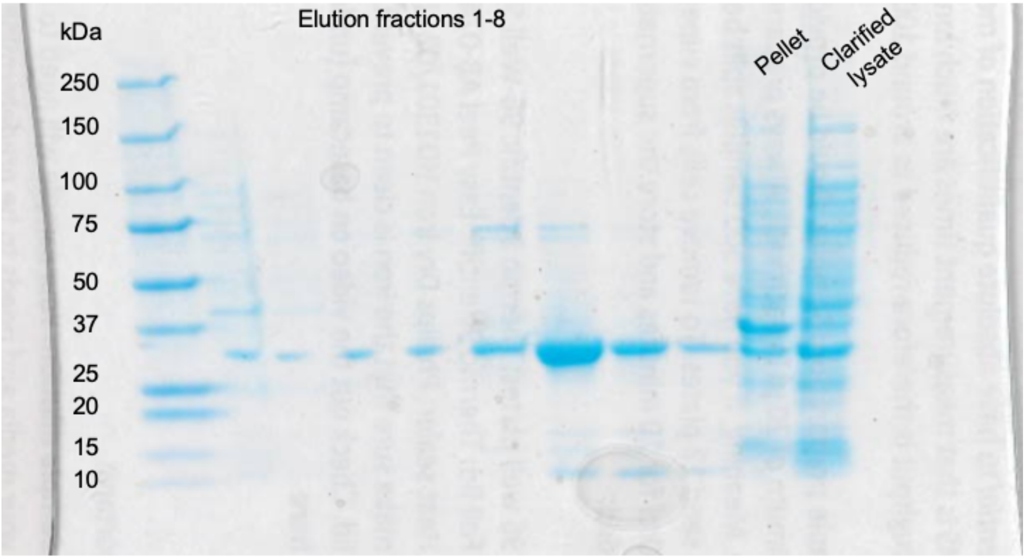


Figure S10. SDS-PAGE gel analysis of purified MBX3358097.1. The protein markers of 250 kDa (Thermo Scientific™ PageRuler™, cropped) is in lane 1. Lanes 2-9 show the elution fractions after expressing the 34 kDa protein.

Table S1. Metagenomes in this study (n=75) from two independent experiments, including name, type, time of collection (D = day) and raw reads per sample.

| experiment_ number | sample_id | raw_reads | sample_type | q20_percent | q30_percent | gc_percent | time_of_sampling |
|---|---|---|---|---|---|---|---|
| 1 | A01 | 121341888 | Biofilm | 98.19 | 94.65 | 52.49 | D29 |
| 1 | A02 | 116365900 | Biofilm | 98.13 | 94.53 | 53.94 | D29 |
| 1 | A03 | 99505308 | Biofilm | 98.21 | 94.69 | 52.74 | D29 |
| 1 | A05 | 81447636 | Biofilm | 98.11 | 94.42 | 48.89 | D29 |
| 1 | A06 | 112998324 | Biofilm | 98.1 | 94.42 | 55.8 | D29 |
| 1 | A07 | 141500210 | Biofilm | 98.15 | 94.55 | 55.39 | D29 |
| 1 | A08 | 145270964 | Biofilm | 98.15 | 94.84 | 57.09 | D29 |
| 1 | A09 | 166698384 | Biofilm | 98.36 | 95.35 | 58.27 | D29 |
| 1 | A10 | 134375116 | Biofilm | 98.24 | 95.09 | 56.27 | D29 |
| 1 | A11 | 134317306 | Biofilm | 98.45 | 95.52 | 56.66 | D29 |
| 1 | A12 | 169624052 | Biofilm | 98.41 | 95.42 | 56.39 | D29 |
| 1 | A13 | 118298608 | Biofilm | 98.29 | 95.21 | 57.16 | D29 |
| 1 | A14 | 59727512 | Biofilm | 97.55 | 93.06 | 56.88 | D29 |

| | | | | | | | |
|---|---|---|---|---|---|---|---|
| 1 | A15 | 142648954 | Biofilm | 98.39 | 95.44 | 57.93 | D29 |
| 1 | A16 | 148978274 | Biofilm | 98.45 | 95.58 | 57.43 | D29 |
| 1 | A17 | 143222686 | Wastewater | 98.32 | 95.15 | 55.47 | D07 |
| 1 | A18 | 168783326 | Wastewater | 98.14 | 94.68 | 56.58 | D14 |
| 1 | A19 | 152857740 | Wastewater | 98.28 | 95.08 | 57.27 | Week3 |
| 1 | A20 | 153580858 | Wastewater | 98.34 | 95.26 | 56.51 | D29 |
| 2 | C141 | 133920098 | Biofilm | 98.15 | 94.83 | 53.63 | D29 |
| 2 | C142 | 163614702 | Biofilm | 98.25 | 95.04 | 51.99 | D29 |
| 2 | C143 | 91250104 | Biofilm | 98.06 | 94.81 | 50.9 | D29 |
| 2 | C144 | 114226966 | Biofilm | 98.33 | 95.3 | 55.76 | D29 |
| 2 | C145 | 129540578 | Biofilm | 98.19 | 95.05 | 55.25 | D29 |
| 2 | C146 | 134812260 | Biofilm | 98.08 | 94.63 | 52.14 | D29 |
| 2 | C147 | 121847210 | Biofilm | 98.26 | 95.21 | 55.02 | D29 |
| 2 | C148 | 139620394 | Biofilm | 98.08 | 94.66 | 53.54 | D29 |
| 2 | C149 | 126141656 | Biofilm | 98 | 94.56 | 52.72 | D29 |
| 2 | C150 | 138396090 | Biofilm | 98.18 | 94.84 | 52.51 | D29 |
| 2 | C151 | 126568992 | Biofilm | 97.95 | 94.34 | 54.17 | D29 |

| | | | | | | | |
|---|---|---|---|---|---|---|---|
| 2 | C152 | 32042116 | Biofilm | 97.65 | 93.72 | 54.02 | D29 |
| 2 | C153 | 20420768 | Biofilm | 97.84 | 94.07 | 52.97 | D29 |
| 2 | C154 | 152244882 | Biofilm | 97.99 | 94.47 | 54.56 | D29 |
| 2 | C155 | 151617768 | Biofilm | 98.09 | 94.72 | 55.3 | D29 |
| 2 | C156 | 178335196 | Biofilm | 98.22 | 95 | 55.43 | D29 |
| 2 | C157 | 9003476 | Biofilm | 97.4 | 93.04 | 52.43 | D29 |
| 2 | C158 | 160873350 | Biofilm | 98.14 | 94.84 | 54.39 | D29 |
| 2 | C159 | 10355276 | Biofilm | 97.68 | 93.69 | 52.39 | D29 |
| 2 | C160 | 136376350 | Biofilm | 98.16 | 94.93 | 54.7 | D29 |
| 2 | C161 | 140524964 | Wastewater | 98.25 | 94.92 | 55.23 | D02 |
| 2 | C162 | 144214308 | Wastewater | 98.4 | 95.26 | 54.32 | D04 |
| 2 | C163 | 132093290 | Wastewater | 98.25 | 95.05 | 56.26 | D07 |
| 2 | C164 | 144224486 | Wastewater | 98.34 | 95.21 | 58.15 | D09 |
| 2 | C165 | 183524464 | Wastewater | 98.35 | 95.34 | 59.21 | D11 |
| 2 | C166 | 179825022 | Wastewater | 98.29 | 95.21 | 59.8 | D14 |
| 2 | C167 | 154372100 | Wastewater | 98.39 | 95.48 | 59 | D16 |
| 2 | C168 | 150617742 | Wastewater | 98.22 | 94.98 | 59.61 | D18 |

| | | | | | | | |
|---|---|---|---|---|---|---|---|
| 2 | C169 | 144066302 | Wastewater | 98.28 | 95.26 | 58.37 | D21 |
| 2 | C170 | 163531286 | Wastewater | 98.14 | 94.86 | 58.32 | D23 |
| 2 | C171 | 142066938 | Wastewater | 98.13 | 94.84 | 58.42 | D25 |
| 2 | C172 | 29773262 | Wastewater | 97.78 | 93.81 | 59.37 | D28 |
| 2 | C173 | 153629740 | River_water | 98.17 | 94.92 | 56.92 | D02 |
| 2 | C174 | 139568188 | River_water | 98.36 | 95.15 | 55.72 | D04 |
| 2 | C175 | 151452306 | River_water | 98.35 | 95.33 | 58.25 | D07 |
| 2 | C176 | 158254166 | River_water | 98.21 | 95.03 | 58.03 | D09 |
| 2 | C177 | 183112680 | River_water | 98.19 | 94.98 | 58.18 | D11 |
| 2 | C178 | 160798168 | River_water | 98.21 | 95.05 | 58.48 | D14 |
| 2 | C179 | 169310540 | River_water | 98.15 | 94.74 | 54.72 | D16 |
| 2 | C180 | 138738880 | River_water | 98.29 | 95.13 | 55.04 | D18 |
| 2 | C181 | 138050702 | River_water | 98.26 | 95.1 | 55.89 | D21 |
| 2 | C182 | 112023898 | River_water | 98.32 | 95.05 | 50.71 | D23 |
| 2 | C183 | 141217696 | River_water | 98.18 | 94.82 | 54.97 | D25 |
| 2 | C184 | 11659254 | River_water | 97.31 | 92.4 | 53.18 | D28 |
| 2 | C197 | 160088318 | Filtrated_wastewater | 98.25 | 95.06 | 60.43 | D02 |

| | | | | | | | |
|---|---|---|---|---|---|---|---|
| 2 | C198 | 156079092 | Filtrated_wastewater | 98.2 | 94.93 | 58.86 | D04 |
| 2 | C199 | 15499038 | Filtrated_wastewater | 98.02 | 94.21 | 48.05 | D07 |
| 2 | C200 | 163073804 | Filtrated_wastewater | 97.6 | 93.03 | 52.77 | D09 |
| 2 | C201 | 163207246 | Filtrated_wastewater | 98.2 | 95 | 57.74 | D11 |
| 2 | C202 | 7824054 | Filtrated_wastewater | 97.71 | 93.81 | 56.73 | D14 |
| 2 | C203 | 182220412 | Filtrated_wastewater | 98.02 | 94.51 | 52.81 | D16 |
| 2 | C204 | 204205456 | Filtrated_wastewater | 98.52 | 95.81 | 65.05 | D18 |
| 2 | C205 | 170904930 | Filtrated_wastewater | 98.31 | 95.32 | 61.35 | D21 |
| 2 | C206 | 181968864 | Filtrated_wastewater | 97.9 | 93.9 | 62.12 | D23 |
| 2 | C207 | 164511632 | Filtrated_wastewater | 98.27 | 95.13 | 61.41 | D25 |
| 2 | C208 | 152281054 | Filtrated_wastewater | 98.33 | 95.3 | 61.14 | D28 |

Table S2. Parameters used for read mapping with bowtie2.

| ***Flag*** | ***Function*** |
|---|---|
| **--local** | Perform local read alignment |
| **-D 20** | 20 seed extension attempts can fail before bowtie2 proceeds with existing alignments |
| **-R 3** | bowtie2 will re-seed reads with repetitive seeds up to 3 times |
| **-L 3** | Sets the length of seed substrings to align during multistring alignment to 3 |
| **-N 1** | Sets the number of mismatches to allowed in a seed alignment during multiseed alignment |
| **--gbar 1** | Disallow gaps within 1 position of the beginning or end of a read |
| **--mp 3** | Sets maximum mismatch penalty to 3 |
| **-p 4** | Use 4 cores for parallelisation |

Table S3. Information about the synthesized *Phycisphaeraceae* beta-lactamase gBlocks.

| | |
|---|---|
| Vector used | pCDF-Duet-1 |
| Cut-sites for insertion | BamHI/HindIII (Multiple cloning site 1, MCS-1) |

| | |
|---|---|
| 6X-His-tag location | N-terminus |
| Protein sequence for MBX3358097.1<br><br>(Signal peptide sequence in bold, removed from synthesized DNA sequence) | **MNQQRRDLLRLTAGLAGLGLVGGPAVAAATASGARVTRR**SGDRDRYFTWEQPG<br>GNDRPIWVAVGEGGNSLVLVGEDESLVVDCKTAAFGAALLREANSLSSRAPVK<br>TVINTHHHADHTGGNYAMIAAGLPVISHAKCAKRVASGENRDRYAPHVEAAIR<br>ALAKSDSETAKAIMAEAESLAAKIQDLPPAAFAPTKTITENTEMEAASERLLL<br>QHAGNGHTDNDVFVFIPRLNVLHTGDLLFARSHPFIDRAAGASTTGWCDAVRR<br>MITLCDSETIVVPGHGDVTDLEGLKAQIVYFDTVRDGVSKMIDAGKSRDDVTR<br>SELEAYADYARPQLRPAAFGAVYDEIIEERQRDEGH |
| Codon-optimized DNA sequence for MBX3358097.1<br><br>(Homology arms for Gibson assembly shown in red) | ACCATCATCACCACAGCCAGTCCGGTGACCGCGACCGCTACTTCACATGGGAA<br>CAGCCAGGAGGTAACGACCGTCCGATTTGGGTGGCCGTTGGCGAAGGGGGAAA<br>TAGTTTAGTTTTGGTGGGTGAGGATGAGTCCTTGGTTGTCGATTGCAAAACTG<br>CTGCCTTCGGCGCGGCGCTTTTACGCGAGGCTAACTCCTTATCGAGCCGCGCC<br>CCCGTAAAGACTGTGATCAATACTCACCACCACGCTGACCATACCGGAGGTAA<br>CTACGCAATGATTGCAGCGGGCTTGCCCGTCATTAGTCATGCCAAGTGCGCCA<br>AACGTGTGGCCAGCGGCGAAAATCGCGACCGCTACGCCCCTCATGTGGAGGCC<br>GCCATCCGTGCCTTGGCAAAGTCCGATTCTGAGACCGCCAAAGCTATTATGGC<br>AGAAGCGGAGAGTCTGGCAGCCAAAATCCAAGATTTACCACCTGCCGCGTTTG<br>CCCCAACCAAAACTATCACCGAGAATACTGAAATGGAGGCGGCGAGTGAGCGT<br>CTTTTGCTTCAACATGCGGGTAATGGGCACACAGACAACGACGTGTTTGTTTT<br>TATCCCTCGCTTAAACGTATTACACACGGGGGACTTGCTGTTCGCGCGTAGTC<br>ATCCTTTCATCGACCGCGCTGCAGGAGCTTCTACCACGGGTTGGTGTGACGCG<br>GTTCGTCGCATGATCACGCTGTGTGATTCCGAAACGATTGTAGTACCGGGGCA<br>CGGGGACGTCACGGATCTTGAGGGGTTGAAGGCTCAAATCGTATATTTCGACA<br>CCGTTCGCGATGGCGTCAGCAAGATGATTGATGCTGGAAATCGCGTGATGAC<br>GTAACGCGCAGCGAGTTGGAGGCATACGCTGACTATGCACGTCCGCAGCTTCG<br>TCCAGCCGCTTTTGGAGCAGTTTACGATGAGATCATTGAGGAACGTCAGCGTG<br>ATGAGGGGCACTAATGCGGCCGCATAATGCTTA |
| Protein sequence for MBX3359331.1<br><br>(No signal peptide detected) | MTTTYSWRLLRAGAFRLDGGSMFGLIPRTVWSRDVPTDDRGRISVQHNCLLLE<br>RAGPAPAPAKGSPPLPSPPSSPRLIVIECGTGDKLDAKSRDLFAMEERSVIDA<br>LHEADCRCEDVEAVVVSHLHFDHAGGLTRLCRGGESPDWTGPASTFTGARGDH<br>GVKVTFPNATVHVQRREWEDAIANRSVMTRTYFPDHLQPIRERLSLTDSPRPF<br>SPGVTPGRDEAPAAPVALRETEIFPGVSVFLTPGHTWGQQAVKFTDTDGQTIV |

| | |
|---|---|
| | FTPDVMPTVNHVGAAYSLAYDVEPYTSMVTRRWFLEEAAAAGWTLVLDHEPGD<br>AVRRVAANGKGWFTLQA |
| Codon-optimized DNA sequence for MBX3359331.1<br><br>(Homology arms for Gibson assembly shown in red) | ACCATCATCACCACAGCCAGATGACCACTACCTATAGCTGGCGTTTGCTTCGT<br>GCGGGCGCATTTCGTCTGGATGGAGGCTCCATGTTTGGGCTGATCCCGCGTAC<br>TGTCTGGTCACGCGACGTACCTACCGACGATCGCGGACGCATCAGTGTCCAGC<br>ATAACTGCCTTTTGCTTGAGCGTGCTGGTCCAGCTCCGGCACCCGCCAAGGGG<br>TCGCCACCACTGCCCTCCCCCCCAAGCTCGCCTCGTTTAATCGTGATCGAATG<br>TGGAACTGGGGACAAGTTGGATGCCAAGTCGCGTGACTTATTTGCGATGGAAG<br>AACGTTCGGTTATTGATGCGTTGCATGAAGCGGACTGTCGTTGCGAAGATGTC<br>GAAGCGGTCGTAGTATCCCATCTGCATTTTGATCATGCGGGAGGATTAACCCG<br>CCTTTGCCGCGGTGGAGAAAGTCCAGATTGGACCGGACCTGCGAGCACTTTCA<br>CTGGAGCCCGTGGTGATCACGGAGTGAAAGTGACCTTTCCTAACGCAACCGTC<br>CACGTTCAACGCCGTGAATGGGAGGACGCGATCGCAAACCGTAGTGTAATGAC<br>ACGTACCTATTTTCCAGACCACCTTCAGCCTATCCGCGAGCGTCTGAGTTTAA<br>CGGACTCGCCGCGCCCCTTTTCGCCCGGGGTTACACCTGGACGCGATGAAGCT<br>CCGGCGGCCCCCGTGGCACTTCGTGAGACCGAGATTTTCCCAGGAGTTTCGGT<br>GTTCTTGACTCCTGGCCATACTTGGGGTCAACAGGCTGTCAAGTTTACAGATA<br>CGGACGGACAAACTATTGTTTTTACGCCTGATGTAATGCCTACAGTGAACCAC<br>GTAGGTGCTGCGTATAGCTTAGCCTACGACGTAGAGCCCTACACCTCAATGGT<br>CACACGTCGCTGGTTTCTGGAGGAGGCCGCGGCAGCAGGTTGGACGCTGGTCT<br>TGGATCATGAGCCTGGTGATGCCGTACGCCGTGTTGCAGCTAACGGGAAAGGT<br>TGGTTCACTTTACAAGCCTGATGCGGCCGCATAATGCTTA |

**Table S4.** Mean concentrations of MP in biofilms normalized by biomass (ng MP/mg AFDW biofilm ± standard error) measured across all WW% summarized from Desiante et al. [11]. The selected group of 27 MPs were retained from the original dataset that consisted of 50 MPs, the selection criteria was discarding the MPs with more than 3 concentrations lower than the quantification limit (LOQ).

* < LOQ = less than the Limit of Quantification

| Substance group | Compound | 00% WW | 30% WW | 30% WW UF | 80% WW | 80% WW UF |
|---|---|---|---|---|---|---|
| *Antibiotics* | Clarithromycin | < LOQ* | 45.91 ± 5.7 | 51.24 ± 10.45 | 94.43 ± 15.31 | 91.15 ± 25.4 |
| | Sulfamethoxazole | < LOQ* | < LOQ* | < LOQ* | 4.46 ± 1.14 | 4.12 ± 0.43 |
| | Trimethoprim | < LOQ* | 7.31 ± 0.97 | 4.91 ± 0.95 | 9.3 ± 0.92 | 6.46 ± 0.87 |
| *Artificial sweeteners* | Acesulfame-K | < LOQ * | 59.2 ± 2.17 | 70.56 ± 6.87 | 90.97 ± 14.95 | 181 ± 16.63 |
| | Cyclamate | 22.95 ± 0.52 | 12.84 ± 1.91 | 20.16 ± 2.55 | 10.56 ± 2.14 | 16.4 ± 3.93 |
| *Corrosion inhibitors* | 4/5-Methylbenzotriazole | 77.51 ± 4.52 | 63.22 ± 3.51 | 66.16 ± 7.35 | 50.68 ± 5.59 | 30.61 ± 4.16 |
| | Benzotriazole | 296.24 ± 60.38 | 398.87 ± 84.61 | 180.26 ± 22.99 | 387.94 ± 64.91 | 770.46 ± 524.87 |
| *Pesticides* | DEET | 32.12 ± 2.4 | 22.52 ± 1 | 21.41 ± 1.28 | 28.79 ± 4.93 | 22.64 ± 1.75 |
| | Diuron | 25.8 ± 5.25 | 52.38 ± 6.86 | 41.11 ± 6.47 | 75.17 ± 5.69 | 58.5 ± 7.29 |
| | Isoproturon | < LOQ* | 2.35 ± 0.26 | 2.2 ± 0.43 | 7.04 ± 2.29 | 2.54 ± 0.35 |
| | Terbutryn | 48.18 ± 4.06 | 118.7 ± 10.48 | 95.15 ± 14.88 | 178.82 ± 3.14 | 198.4 ± 12.15 |

| | | | | | | |
|---|---|---|---|---|---|---|
| *Pharmaceuticals* | 4-Acetamidoantipyrine | < LOQ* | 9.99 ± 1.51 | 5.36 ± 0.5 | 10.26 ± 1.62 | 12.19 ± 1.41 |
| | 4-Formylaminoantipyrine | < LOQ* | < LOQ* | < LOQ* | 42.29 ± 6.19 | 36.16 ± 0.63 |
| | Amisulpride | 0.9 ± 0.09 | 4.3 ± 0.79 | 4.23 ± 0.59 | 5.58 ± 0.98 | 3.81 ± 0.45 |
| | Atenolol | < LOQ* | < LOQ* | < LOQ* | 3.92 ± 0.24 | 4.38 ± 0.14 |
| | Carbamazepine | < LOQ* | 4.25 ± 0.24 | 2.73 ± 0.3 | 10.02 ± 1.36 | 6.72 ± 0.72 |
| | Cetirizine | < LOQ* | 8.12 ± 0.19 | < LOQ* | 14.41 ± 5.08 | 6.4 |
| | Clopidogrel-Carboxylic-Acid | < LOQ* | < LOQ* | < LOQ* | 5.21 ± 1.03 | 3.73 ± 0.32 |
| | Diclofenac | < LOQ* | 34.62 ± 4.59 | 19.93 ± 1.2 | 64.72 ± 6.35 | 67.82 ± 14.42 |
| | Hydrochlorothiazide | < LOQ* | 14.09 ± 1.22 | 12.75 | 36.33 ± 4.34 | 29.53 ± 4.38 |
| | Lamotrigine | 29.22 ± 3.59 | 38.37 ± 2.63 | 38.17 ± 6.12 | 56.55 ± 8.5 | 50.3 ± 6.22 |
| | Lidocaine (Diocaine) | < LOQ* | 17.58 ± 1.83 | 12.81 ± 1.35 | 40.72 ± 5.38 | 28.55 ± 3.22 |
| | Mefenamic acid | < LOQ* | 16.17 ± 1.55 | 5.66 ± 0.69 | 18.05 ± 2.25 | 16.7 ± 2.96 |
| | Metoprolol | 3.5 ± 0.54 | 25.07 ± 2.38 | 23.89 ± 2.83 | 54.67 ± 5.77 | 71 ± 8.42 |

| | | | | | | |
|---|---|---|---|---|---|---|
| | Sitagliptin | 15.82 ± 1.24 | 31.96 ± 2.62 | 40.93 ± 4.74 | 44.86 ± 7.33 | 44.96 ± 3.94 |
| | Venlafaxine | 3.2 ± 0.5 | 59.24 ± 5.54 | 45.21 ± 6.77 | 168.6 ± 17.99 | 133.02 ± 17.15 |
| *Tracer* | Caffeine | 67.48 ± 29.51 | 82.35 ± 55.84 | 46.51 ± 10.33 | 168.09 ± 90.22 | 148 ± 104.24 |

Table S5. ARGs with highly significant differences (Kruskal-Wallis test + Benjamini-Hochberg p-value correction) in biofilm across all conditions.

| **Sample** | **Sample type** | **Treatment** | **ARG class** | **Read count** | **Adjusted P-value** |
|---|---|---|---|---|---|
| C141 | Biofilm | WW30 | aminoglycoside | 0.023650528 | 0.0000022036 |
| C141 | Biofilm | WW30 | beta_lactam | 0.028009139 | 0.0000001720 |
| C141 | Biofilm | WW30 | fluoroquinolone | 0.006167367 | 0.0001397333 |
| C141 | Biofilm | WW30 | glycopeptide | 0.002955254 | 0.0000003264 |
| C141 | Biofilm | WW30 | mls | 0.040964759 | 0.0004164706 |
| C141 | Biofilm | WW30 | multidrug | 0.893743148 | 0.0000004573 |

| | | | | | |
|---|---|---|---|---|---|
| C141 | Biofilm | WW30 | peptide | 0.014213929 | 0.0000001720 |
| C141 | Biofilm | WW30 | rifamycin | 0.047012975 | 0.0000021400 |
| C141 | Biofilm | WW30 | sulfonamide | 0.001266985 | 0.0000022036 |
| C141 | Biofilm | WW30 | tetracycline | 0.012780879 | 0.0000000295 |
| C142 | Biofilm | WW80 | aminoglycoside | 0.025250839 | 0.0000022036 |
| C142 | Biofilm | WW80 | beta_lactam | 0.058679181 | 0.0000001720 |
| C142 | Biofilm | WW80 | fluoroquinolone | 0.005105337 | 0.0001397333 |
| C142 | Biofilm | WW80 | glycopeptide | 0.00308333 | 0.0000003264 |
| C142 | Biofilm | WW80 | mls | 0.025152491 | 0.0004164706 |
| C142 | Biofilm | WW80 | multidrug | 0.771270694 | 0.0000004573 |
| C142 | Biofilm | WW80 | peptide | 0.019917616 | 0.0000001720 |
| C142 | Biofilm | WW80 | rifamycin | 0.043394096 | 0.0000021400 |

| | | | | | |
|---|---|---|---|---|---|
| C142 | Biofilm | WW80 | sulfonamide | 0.0007827 | 0.0000022036 |
| C142 | Biofilm | WW80 | tetracycline | 0.010350101 | 0.0000000295 |
| C143 | Biofilm | WW80UF | aminoglycoside | 0.027046465 | 0.0000022036 |
| C143 | Biofilm | WW80UF | beta_lactam | 0.00417183 | 0.0000001720 |
| C143 | Biofilm | WW80UF | fluoroquinolone | 0.002547184 | 0.0001397333 |
| C143 | Biofilm | WW80UF | glycopeptide | 0.000800767 | 0.0000003264 |
| C143 | Biofilm | WW80UF | mls | 0.037897519 | 0.0004164706 |
| C143 | Biofilm | WW80UF | multidrug | 0.67735157 | 0.0000004573 |
| C143 | Biofilm | WW80UF | peptide | 0.008051203 | 0.0000001720 |
| C143 | Biofilm | WW80UF | rifamycin | 0.006565144 | 0.0000021400 |
| C143 | Biofilm | WW80UF | sulfonamide | 0.003794449 | 0.0000022036 |
| C143 | Biofilm | WW80UF | tetracycline | 0.002466378 | 0.0000000295 |

| | | | | | |
|---|---|---|---|---|---|
| C144 | Biofilm | WW00 | aminoglycoside | 0.024541346 | 0.0000022036 |
| C144 | Biofilm | WW00 | beta_lactam | 0.004318396 | 0.0000001720 |
| C144 | Biofilm | WW00 | fluoroquinolone | 0.002275699 | 0.0001397333 |
| C144 | Biofilm | WW00 | glycopeptide | 0.002151262 | 0.0000003264 |
| C144 | Biofilm | WW00 | mls | 0.018292163 | 0.0004164706 |
| C144 | Biofilm | WW00 | multidrug | 0.732490251 | 0.0000004573 |
| C144 | Biofilm | WW00 | peptide | 0.004865103 | 0.0000001720 |
| C144 | Biofilm | WW00 | rifamycin | 0.015057834 | 0.0000021400 |
| C144 | Biofilm | WW00 | sulfonamide | 0.001106303 | 0.0000022036 |
| C144 | Biofilm | WW00 | tetracycline | 0.002843233 | 0.0000000295 |
| C145 | Biofilm | WW30UF | aminoglycoside | 0.023547365 | 0.0000022036 |
| C145 | Biofilm | WW30UF | beta_lactam | 0.002200755 | 0.0000001720 |

| | | | | | |
|---|---|---|---|---|---|
| C145 | Biofilm | WW30UF | fluoroquinolone | 0.001763619 | 0.0001397333 |
| C145 | Biofilm | WW30UF | glycopeptide | 0.003456219 | 0.0000003264 |
| C145 | Biofilm | WW30UF | mls | 0.03142258 | 0.0004164706 |
| C145 | Biofilm | WW30UF | multidrug | 0.783664287 | 0.0000004573 |
| C145 | Biofilm | WW30UF | peptide | 0.004275782 | 0.0000001720 |
| C145 | Biofilm | WW30UF | rifamycin | 0.015622466 | 0.0000021400 |
| C145 | Biofilm | WW30UF | sulfonamide | 0.001516421 | 0.0000022036 |
| C145 | Biofilm | WW30UF | tetracycline | 0.003934724 | 0.0000000295 |
| C146 | Biofilm | WW30 | aminoglycoside | 0.019316387 | 0.0000022036 |
| C146 | Biofilm | WW30 | beta_lactam | 0.031785726 | 0.0000001720 |
| C146 | Biofilm | WW30 | fluoroquinolone | 0.003711616 | 0.0001397333 |
| C146 | Biofilm | WW30 | glycopeptide | 0.003126446 | 0.0000003264 |

| | | | | | |
|---|---|---|---|---|---|
| C146 | Biofilm | WW30 | mls | 0.026157307 | 0.0004164706 |
| C146 | Biofilm | WW30 | multidrug | 0.769512276 | 0.0000004573 |
| C146 | Biofilm | WW30 | peptide | 0.011350981 | 0.0000001720 |
| C146 | Biofilm | WW30 | rifamycin | 0.035200721 | 0.0000021400 |
| C146 | Biofilm | WW30 | sulfonamide | 0.001763656 | 0.0000022036 |
| C146 | Biofilm | WW30 | tetracycline | 0.010104327 | 0.0000000295 |
| C147 | Biofilm | WW00 | aminoglycoside | 0.023163666 | 0.0000022036 |
| C147 | Biofilm | WW00 | beta_lactam | 0.003028062 | 0.0000001720 |
| C147 | Biofilm | WW00 | fluoroquinolone | 0.001907805 | 0.0001397333 |
| C147 | Biofilm | WW00 | glycopeptide | 0.0040111 | 0.0000003264 |
| C147 | Biofilm | WW00 | mls | 0.020610744 | 0.0004164706 |
| C147 | Biofilm | WW00 | multidrug | 0.634085484 | 0.0000004573 |

| | | | | | |
|---|---|---|---|---|---|
| C147 | Biofilm | WW00 | peptide | 0.004334097 | 0.0000001720 |
| C147 | Biofilm | WW00 | rifamycin | 0.011661342 | 0.0000021400 |
| C147 | Biofilm | WW00 | sulfonamide | 0.001330959 | 0.0000022036 |
| C147 | Biofilm | WW00 | tetracycline | 0.00238359 | 0.0000000295 |
| C148 | Biofilm | WW80UF | aminoglycoside | 0.011592417 | 0.0000022036 |
| C148 | Biofilm | WW80UF | beta_lactam | 0.003939249 | 0.0000001720 |
| C148 | Biofilm | WW80UF | fluoroquinolone | 0.002069232 | 0.0001397333 |
| C148 | Biofilm | WW80UF | glycopeptide | 0.001407513 | 0.0000003264 |
| C148 | Biofilm | WW80UF | mls | 0.027431474 | 0.0004164706 |
| C148 | Biofilm | WW80UF | multidrug | 0.71876293 | 0.0000004573 |
| C148 | Biofilm | WW80UF | peptide | 0.007975216 | 0.0000001720 |
| C148 | Biofilm | WW80UF | rifamycin | 0.008593237 | 0.0000021400 |

| | | | | | |
|---|---|---|---|---|---|
| C148 | Biofilm | WW80UF | sulfonamide | 0.003186858 | 0.0000022036 |
| C148 | Biofilm | WW80UF | tetracycline | 0.00212905 | 0.0000000295 |
| C149 | Biofilm | WW80UF | aminoglycoside | 0.018104165 | 0.0000022036 |
| C149 | Biofilm | WW80UF | beta_lactam | 0.00285953 | 0.0000001720 |
| C149 | Biofilm | WW80UF | fluoroquinolone | 0.001454947 | 0.0001397333 |
| C149 | Biofilm | WW80UF | glycopeptide | 0.000693025 | 0.0000003264 |
| C149 | Biofilm | WW80UF | mls | 0.031937088 | 0.0004164706 |
| C149 | Biofilm | WW80UF | multidrug | 0.555638165 | 0.0000004573 |
| C149 | Biofilm | WW80UF | peptide | 0.003935992 | 0.0000001720 |
| C149 | Biofilm | WW80UF | rifamycin | 0.010869557 | 0.0000021400 |
| C149 | Biofilm | WW80UF | sulfonamide | 0.002189275 | 0.0000022036 |
| C149 | Biofilm | WW80UF | tetracycline | 0.00159057 | 0.0000000295 |

| | | | | | |
|---|---|---|---|---|---|
| C150 | Biofilm | WW80 | aminoglycoside | 0.033321721 | 0.0000022036 |
| C150 | Biofilm | WW80 | beta_lactam | 0.065553718 | 0.0000001720 |
| C150 | Biofilm | WW80 | fluoroquinolone | 0.008969699 | 0.0001397333 |
| C150 | Biofilm | WW80 | glycopeptide | 0.005405615 | 0.0000003264 |
| C150 | Biofilm | WW80 | mls | 0.02231054 | 0.0004164706 |
| C150 | Biofilm | WW80 | multidrug | 1.295122962 | 0.0000004573 |
| C150 | Biofilm | WW80 | peptide | 0.022838382 | 0.0000001720 |
| C150 | Biofilm | WW80 | rifamycin | 0.088060495 | 0.0000021400 |
| C150 | Biofilm | WW80 | sulfonamide | 0.002552657 | 0.0000022036 |
| C150 | Biofilm | WW80 | tetracycline | 0.021012451 | 0.0000000295 |
| C151 | Biofilm | WW30 | aminoglycoside | 0.044443879 | 0.0000022036 |
| C151 | Biofilm | WW30 | beta_lactam | 0.039519665 | 0.0000001720 |

| | | | | | |
|---|---|---|---|---|---|
| C151 | Biofilm | WW30 | fluoroquinolone | 0.006525304 | 0.0001397333 |
| C151 | Biofilm | WW30 | glycopeptide | 0.004206528 | 0.0000003264 |
| C151 | Biofilm | WW30 | mls | 0.029269674 | 0.0004164706 |
| C151 | Biofilm | WW30 | multidrug | 1.238590717 | 0.0000004573 |
| C151 | Biofilm | WW30 | peptide | 0.01957089 | 0.0000001720 |
| C151 | Biofilm | WW30 | rifamycin | 0.036269244 | 0.0000021400 |
| C151 | Biofilm | WW30 | sulfonamide | 0.002497839 | 0.0000022036 |
| C151 | Biofilm | WW30 | tetracycline | 0.012811294 | 0.0000000295 |
| C152 | Biofilm | WW30UF | aminoglycoside | 0.028928592 | 0.0000022036 |
| C152 | Biofilm | WW30UF | beta_lactam | 0.003780003 | 0.0000001720 |
| C152 | Biofilm | WW30UF | fluoroquinolone | 0.003328769 | 0.0001397333 |
| C152 | Biofilm | WW30UF | glycopeptide | 0.001812082 | 0.0000003264 |

| | | | | | |
|---|---|---|---|---|---|
| C152 | Biofilm | WW30UF | mls | 0.025897977 | 0.0004164706 |
| C152 | Biofilm | WW30UF | multidrug | 0.792237395 | 0.0000004573 |
| C152 | Biofilm | WW30UF | peptide | 0.002585108 | 0.0000001720 |
| C152 | Biofilm | WW30UF | rifamycin | 0.014921063 | 0.0000021400 |
| C152 | Biofilm | WW30UF | sulfonamide | 0.003148412 | 0.0000022036 |
| C152 | Biofilm | WW30UF | tetracycline | 0.003564795 | 0.0000000295 |
| C153 | Biofilm | WW80 | aminoglycoside | 0.02521926 | 0.0000022036 |
| C153 | Biofilm | WW80 | beta_lactam | 0.127869288 | 0.0000001720 |
| C153 | Biofilm | WW80 | fluoroquinolone | 0.013576495 | 0.0001397333 |
| C153 | Biofilm | WW80 | glycopeptide | 0.004564804 | 0.0000003264 |
| C153 | Biofilm | WW80 | mls | 0.03495245 | 0.0004164706 |
| C153 | Biofilm | WW80 | multidrug | 1.676226045 | 0.0000004573 |

| | | | | | |
|---|---|---|---|---|---|
| C153 | Biofilm | WW80 | peptide | 0.038390237 | 0.0000001720 |
| C153 | Biofilm | WW80 | rifamycin | 0.109182903 | 0.0000021400 |
| C153 | Biofilm | WW80 | sulfonamide | 0.004326079 | 0.0000022036 |
| C153 | Biofilm | WW80 | tetracycline | 0.025443481 | 0.0000000295 |
| C154 | Biofilm | WW30UF | aminoglycoside | 0.020641248 | 0.0000022036 |
| C154 | Biofilm | WW30UF | beta_lactam | 0.003950204 | 0.0000001720 |
| C154 | Biofilm | WW30UF | fluoroquinolone | 0.002973907 | 0.0001397333 |
| C154 | Biofilm | WW30UF | glycopeptide | 0.003365285 | 0.0000003264 |
| C154 | Biofilm | WW30UF | mls | 0.013137556 | 0.0004164706 |
| C154 | Biofilm | WW30UF | multidrug | 0.753229264 | 0.0000004573 |
| C154 | Biofilm | WW30UF | peptide | 0.005353975 | 0.0000001720 |
| C154 | Biofilm | WW30UF | rifamycin | 0.011495949 | 0.0000021400 |

| | | | | | |
|---|---|---|---|---|---|
| C154 | Biofilm | WW30UF | sulfonamide | 0.001083822 | 0.0000022036 |
| C154 | Biofilm | WW30UF | tetracycline | 0.002147529 | 0.0000000295 |
| C155 | Biofilm | WW00 | aminoglycoside | 0.024685668 | 0.0000022036 |
| C155 | Biofilm | WW00 | beta_lactam | 0.005695968 | 0.0000001720 |
| C155 | Biofilm | WW00 | fluoroquinolone | 0.002013704 | 0.0001397333 |
| C155 | Biofilm | WW00 | glycopeptide | 0.003836699 | 0.0000003264 |
| C155 | Biofilm | WW00 | mls | 0.043920842 | 0.0004164706 |
| C155 | Biofilm | WW00 | multidrug | 0.864773363 | 0.0000004573 |
| C155 | Biofilm | WW00 | peptide | 0.007158915 | 0.0000001720 |
| C155 | Biofilm | WW00 | rifamycin | 0.01295781 | 0.0000021400 |
| C155 | Biofilm | WW00 | sulfonamide | 0.002424034 | 0.0000022036 |
| C155 | Biofilm | WW00 | tetracycline | 0.004263051 | 0.0000000295 |

| | | | | | |
|---|---|---|---|---|---|
| C156 | Biofilm | WW80UF | aminoglycoside | 0.015648584 | 0.0000022036 |
| C156 | Biofilm | WW80UF | beta_lactam | 0.006795527 | 0.0000001720 |
| C156 | Biofilm | WW80UF | fluoroquinolone | 0.003132731 | 0.0001397333 |
| C156 | Biofilm | WW80UF | glycopeptide | 0.001132537 | 0.0000003264 |
| C156 | Biofilm | WW80UF | mls | 0.014612333 | 0.0004164706 |
| C156 | Biofilm | WW80UF | multidrug | 0.978402284 | 0.0000004573 |
| C156 | Biofilm | WW80UF | peptide | 0.005402871 | 0.0000001720 |
| C156 | Biofilm | WW80UF | rifamycin | 0.01308217 | 0.0000021400 |
| C156 | Biofilm | WW80UF | sulfonamide | 0.004254557 | 0.0000022036 |
| C156 | Biofilm | WW80UF | tetracycline | 0.003445657 | 0.0000000295 |
| C157 | Biofilm | WW30 | aminoglycoside | 0.016717143 | 0.0000022036 |
| C157 | Biofilm | WW30 | beta_lactam | 0.063818426 | 0.0000001720 |

| | | | | | |
|---|---|---|---|---|---|
| C157 | Biofilm | WW30 | fluoroquinolone | 0.029526382 | 0.0001397333 |
| C157 | Biofilm | WW30 | glycopeptide | 0.0029064 | 0.0000003264 |
| C157 | Biofilm | WW30 | mls | 0.005765507 | 0.0004164706 |
| C157 | Biofilm | WW30 | multidrug | 0.992889345 | 0.0000004573 |
| C157 | Biofilm | WW30 | peptide | 0.03180935 | 0.0000001720 |
| C157 | Biofilm | WW30 | rifamycin | 0.04558459 | 0.0000021400 |
| C157 | Biofilm | WW30 | sulfonamide | 0.001147669 | 0.0000022036 |
| C157 | Biofilm | WW30 | tetracycline | 0.015150563 | 0.0000000295 |
| C158 | Biofilm | WW00 | aminoglycoside | 0.021310247 | 0.0000022036 |
| C158 | Biofilm | WW00 | beta_lactam | 0.005032723 | 0.0000001720 |
| C158 | Biofilm | WW00 | fluoroquinolone | 0.001941615 | 0.0001397333 |
| C158 | Biofilm | WW00 | glycopeptide | 0.002014722 | 0.0000003264 |

| | | | | | |
|---|---|---|---|---|---|
| C158 | Biofilm | WW00 | mls | 0.035064192 | 0.0004164706 |
| C158 | Biofilm | WW00 | multidrug | 0.7374293 | 0.0000004573 |
| C158 | Biofilm | WW00 | peptide | 0.00471596 | 0.0000001720 |
| C158 | Biofilm | WW00 | rifamycin | 0.013523442 | 0.0000021400 |
| C158 | Biofilm | WW00 | sulfonamide | 0.000701177 | 0.0000022036 |
| C158 | Biofilm | WW00 | tetracycline | 0.003358845 | 0.0000000295 |
| C159 | Biofilm | WW80 | aminoglycoside | 0.011119977 | 0.0000022036 |
| C159 | Biofilm | WW80 | beta_lactam | 0.101441273 | 0.0000001720 |
| C159 | Biofilm | WW80 | fluoroquinolone | 0.015194762 | 0.0001397333 |
| C159 | Biofilm | WW80 | glycopeptide | 0.001654314 | 0.0000003264 |
| C159 | Biofilm | WW80 | mls | 0.01525623 | 0.0004164706 |
| C159 | Biofilm | WW80 | multidrug | 1.30381725 | 0.0000004573 |

| | | | | | |
|---|---|---|---|---|---|
| C159 | Biofilm | WW80 | peptide | 0.033352765 | 0.0000001720 |
| C159 | Biofilm | WW80 | rifamycin | 0.100543121 | 0.0000021400 |
| C159 | Biofilm | WW80 | sulfonamide | 0.001364351 | 0.0000022036 |
| C159 | Biofilm | WW80 | tetracycline | 0.02983226 | 0.0000000295 |
| C160 | Biofilm | WW30UF | aminoglycoside | 0.025536119 | 0.0000022036 |
| C160 | Biofilm | WW30UF | beta_lactam | 0.004662316 | 0.0000001720 |
| C160 | Biofilm | WW30UF | fluoroquinolone | 0.002975187 | 0.0001397333 |
| C160 | Biofilm | WW30UF | glycopeptide | 0.002312792 | 0.0000003264 |
| C160 | Biofilm | WW30UF | mls | 0.033253452 | 0.0004164706 |
| C160 | Biofilm | WW30UF | multidrug | 0.925972974 | 0.0000004573 |
| C160 | Biofilm | WW30UF | peptide | 0.005558759 | 0.0000001720 |
| C160 | Biofilm | WW30UF | rifamycin | 0.018137157 | 0.0000021400 |

| | | | | | |
|---|---|---|---|---|---|
| C160 | Biofilm | WW30UF | sulfonamide | 0.001790719 | 0.0000022036 |
| C160 | Biofilm | WW30UF | tetracycline | 0.009384694 | 0.0000000295 |

Table S6. ARGs with significant differences in coverage in all conditions according to results of the Kruskal-Wallis test for the normalized average coverage in all (WW%) treatments.

| **Gene ID** | **Corrected p-value** | **Gene type** |
|---|---|---|
| k121_858589_5 | 0.0054 | Class-A beta-lactamase |
| k121_252136_5 | 0.0054 | Class-D beta-lactamase (OXA-4) |
| k121_895379_1 | 0.0334 | Other |
| k121_1259309_2 | 0.0416 | Other |
| k121_671215_2 | 0.0416 | Macrolide-Lincomide-Streptogramin (MLS) |
| k121_1519353_3 | 0.0416 | Other |

| | | |
|---|---|---|
| k121_906279_2 | 0.0467 | Multidrug |
| k121_68825_2 | 0.0487 | Aminoglycoside |
| k121_660679_2 | 0.0488 | Other |
| k121_1269806_2 | 0.0488 | Extended spectrum beta-lactamase BEL-2 |
